# The interplay between peptides and RNA is critical for protoribosome compartmentalization and stability

**DOI:** 10.1101/2024.01.29.577748

**Authors:** Simone Codispoti, Tomoko Yamaguchi, Mikhail Makarov, Valerio G. Giacobelli, Martin Mašek, Michal H. Kolář, Alma Carolina Sanchez Rocha, Kosuke Fujishima, Giuliano Zanchetta, Klára Hlouchová

**Affiliations:** Dipartimento di Biotecnologie Mediche e Medicina Traslazionale, Università di Milano, Segrate, 20054, Italy; Earth-Life Science Institute, Tokyo Institute of Technology, Ookayama, Meguro-ku, Tokyo 152-8550, Japan; Department of Cell Biology, Faculty of Science, Charles University, BIOCEV, Prague, 12843, Czech Republic; Department of Physical Chemistry, University of Chemistry and Technology, Technicka 5, 16628 Prague, Czech Republic; School of Life Science and Technology, Tokyo Institute of Technology, Meguro-ku, Tokyo 152-8550, Japan; Graduate School of Media and Governance, Keio University, Fujisawa, 252-0882, Japan; Institute of Organic Chemistry and Biochemistry, Czech Academy of Sciences, Prague, 16610, Czech Republic

## Abstract

The ribosome, owing to its exceptional conservation and biological importance, harbours a remarkable molecular fossil known as the protoribosome. It surrounds the peptidyl transferase center (PTC), responsible for peptide bond formation. While previous studies have demonstrated the PTC activity in RNA alone, our investigation reveals the intricate roles of the ribosomal protein fragments (rPeptides) within the ribosomal core.

This research highlights the significance of rPeptides in stability and coacervation of two distinct protoribosomal evolutionary stages. The 617nt “big” protoribosome model which associates with rPeptides specifically, exhibits a structurally defined and rigid nature, further stabilised by the peptides. In contrast, the 136nt “small” model, previously linked to peptidyltransferase activity, displays greater structural flexibility. While this construct interacts with rPeptides with lower specificity, they induce coacervation of the “small” protoribosome across a wide concentration range, which is concomitantly dependent on the RNA sequence and structure. Moreover, these conditions protect RNA from degradation. This phenomenon suggests a significant evolutionary advantage in the RNA-protein interaction at the early stages of ribosome evolution.

The distinct properties of the two protoribosomal stages suggest that rPeptides initially provided compartmentalization and prevented RNA degradation, preceding the emergence of specific RNA-protein interactions crucial for the ribosomal structural integrity.

**keypoints:**

- The most ancient fragments of ribosomal peptides trigger coacervation of protoribosome
- The protoribosome coacervation provides protection against RNA degradation
- Coacervation is more profound with a smaller and more flexible model of the protoribosome

## Introduction

Every cell capable of protein expression contains ribosomes. Due to their omnipresence and high conservation across all life forms, ribosomes qualify as ancient molecular fossils, attracting evolutionary biologists as the best connection to our biological past (1) (2). According to several studies, the peptidyl transferase center (PTC) of the ribosome evolved around 3.8-4.2 billion years ago, before the appearance of the last universal common ancestor. As a nest of RNA capable of polymerizing amino acids, the PTC (also known as protoribosome) is considered by some scientists to be a true cradle of life embedded in the ribosome, bridging prebiotic chemistry and protein-dominated biology (3). Although the PTC is made up of ribosomal RNA (rRNA), it is surrounded by fragments of ribosomal proteins (rProteins) that lack secondary-structure motifs and are considered older than rProteins with globular structures found in the outer layers of the modern ribosome (4) (5). A record of polypeptide interaction with the PTC can be found in these inner tails. With the onset of templated synthesis, the sequence of these polypeptides is presumed to have been fixed. They are considered to have expanded into proteins, along with the rRNA, following an accretion process. This is known as the onion model for ribosome evolution (5).

Much of the progress in the study of the protoribosome has resulted from the explosion of resolved ribosomal structures over the last two decades (reflected also in the 2009 Nobel Prize in Chemistry awarded to V. Ramakrishnan, T. A. Steitz, and A. E. Yonath). Based on these, several groups have proposed regions of the 23S rRNA in the large subunit, ranging approximately 100-600 nucleotides (nt) in length, as the ancestral PTC (3) (6) (7) (8) (9). Most importantly, two independent groups have recently demonstrated the peptidyl-transferase activity of their ∼70-140 nt rRNA constructs, mimicking the semi-symmetrical PTC pocket (10) (11) (12). Unlike some of the larger PTC constructs, these minimal protoribosome models were devoid of peptides, proving the rRNA sovereignty in the catalysis of amino acid polymerization.

Although some scientists consider peptides relevant only after the evolution of templated protein synthesis, peptides have been repeatedly reported prebiotically plausible, supported by the abundance of amino acids and the facile nature of their condensation reaction (13). Some of the ribosomal protein fragments/tails (rPeptides) have been implied to interact with a larger model of the PTC and form a specific RNA-peptide assembly (9). Additionally, ribosomal peptides have been reported to enhance RNA polymerase ribozyme function (14).

Here we report the key role of ancient rPeptides in protoribosome stabilisation and coacervation. These prebiotically important properties depend on the RNA length and secondary structure content but they are predominantly influenced by the interplay between RNA and peptides. In this context, we propose a scenario in which peptides facilitated protoribosome accumulation within liquid droplets, thereby fostering the coevolution of a compartmentalised core for an active RNA-peptide world with a prospect of a true protocell, as proposed by Aleksander Oparin a whole century ago (15) (16).

## Material & Methods

### Synthesis of PTC rRNAs and peptides

We adopted two published models of the PTC from *Thermus Thermophilus*, namely WT bPTC (referring to a “big” 617 nt rRNA construct) and WT sPTC (referring to a “small” 136 nt rRNA construct) (9) (10) (11) (**Figure 1**). In both cases, the constructs stem from segmented fragments of the 23S rRNA, connected with loops to preserve the three-dimensional structures. WT bPTC is made up of the 505 nt from the 23S rRNA, joined by 11 stem loops (5’-gccGUAAggc-3’); WT sPTC is composed of 116 nt from the 23S rRNA and connected with 3 stem loops (5’-CUUCGG-3’) (**Supplementary Fig. 1**). As a control, shuffled versions of both PTC rRNAs were designed by randomising their sequences while keeping the same composition of A, U, G and C (**Supplementary Fig. 2**). WT_bPTC and Sh_bPTC ssDNA oligonucleotides were purchased from Integrated DNA Technologies (IDT), and the remaining ssDNA oligos were purchased from Sigma Aldrich (**Supplementary Table 1**).

**Figure 1:**
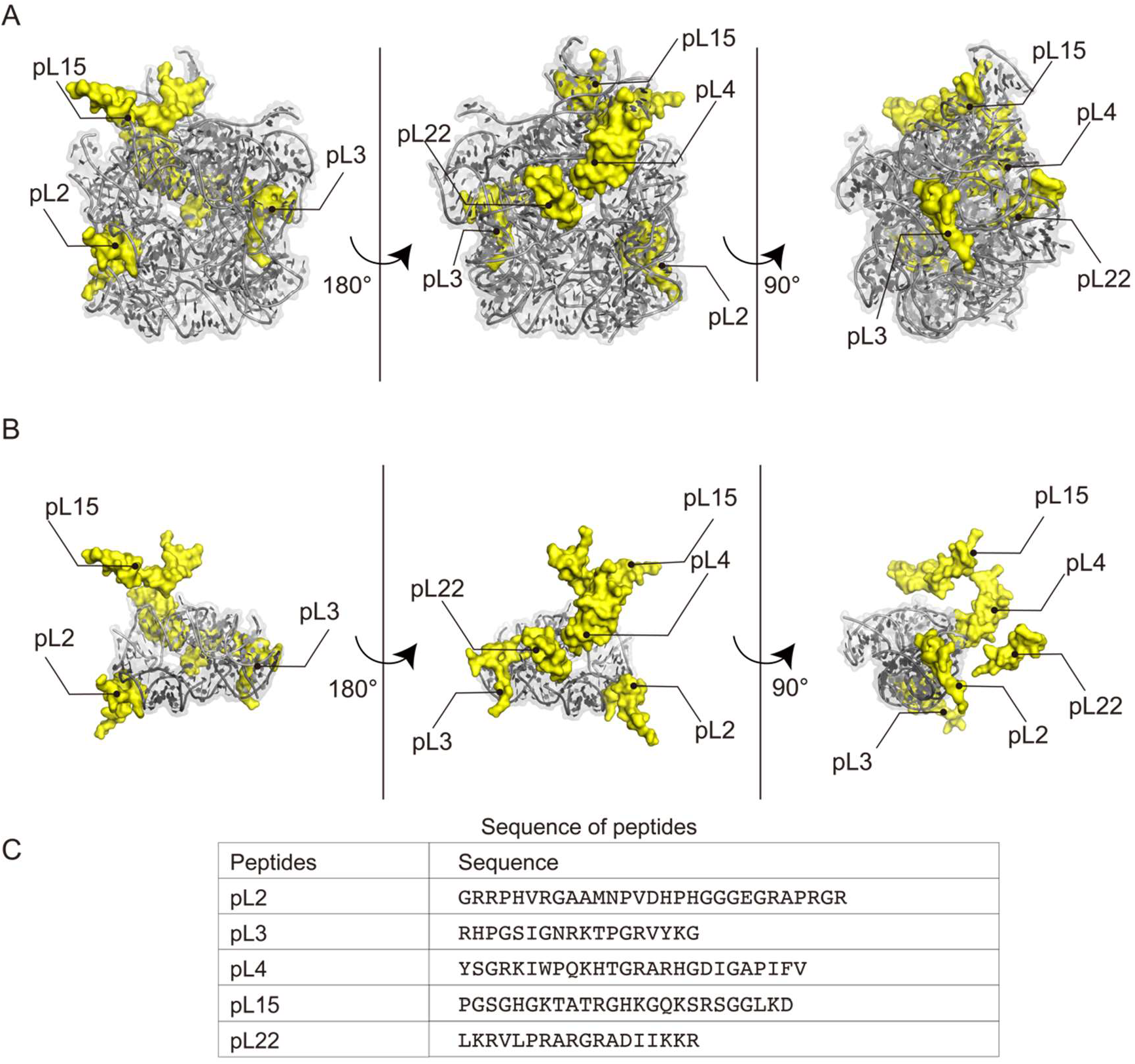
PTC models and rPeptide sequences. (A-B) The models of WT bPTC (*A*) and WT sPTC (*B*) in grey and rPeptides (pL2, 3, 4, 15, 22) in yellow. Both models are extracted from the *Thermus Thermophilus* ribosomal structure (PDB ID: 4V51), missing the loops that were added by authors of the designed models (9) (11). The two symmetrical A and P regions of the WT sPTC (*B*) are depicted with lighter and darker shades of grey, respectively. (C) Sequences of rPeptides used in this study.

For the synthesis of WT and Sh bPTC rRNAs, dsDNA templates were prepared by PCR amplification of WT_bPTC and Sh_bPTC ssDNA oligos using Q5 Hot Start High-Fidelity DNA Polymerase (NEB) according to the manufacturer’s instructions. PCR amplification was performed using bPTC_F and bPTC_R primers according to the following program: initial denaturation at 98°C for 30 s; denaturation at 98°C for 10 s; annealing at 69°C for 30 s; extension at 72°C for 60 s; final extension at 72°C for 20 s; 32 cycles.

For the synthesis of WT sPTC rRNA, dsDNA template was prepared by annealing of WT_sPTC_F and WT_sPTC_R ssDNA oligos (**Supplementary Table 1**) followed by DNA synthesis using Klenow Fragment of DNA polymerase I (NEB). For annealing, 15 pmol of WT_sPTC_F and WT_sPTC_R ssDNA oligos (8.1 μg of DNA in total) were mixed in NEB2 buffer to the total volume of 47 μL, denatured at 80°C for 5 min and cooled down to room temperature at 0.1°C/s rate. Then, 1 μL of 5U/ μL Klenow Fragment of DNA polymerase I (NEB) and 2 μL of 10 mM dNTP mix (SERVA) were added to the reaction mixture, and the reaction mixture was kept at 25°C for 1 hour followed by heat inactivation of enzyme at 50°C for 15 min. For the synthesis of Sh1 and Sh2 sPTC rRNAs, 40 pmol of Sh1_sPTC_R / Sh2_sPTC_R ssDNA and T7_F (**Supplementary Table 1**) was mixed in H_2_O to the total volume of 36 μl, incubated 95°C for 1 min and then kept at room temperature for 2 min; this was used as DNA template (1.9 μg of DNA in total).

The resulting dsDNA templates were purified from 2% TAE-agarose gel using Monarch Gel Dissolving Buffer (NEB) and Monarch PCR&DNA Cleanup Kit (5 μg) (NEB). The purified dsDNA templates were transcribed using HiScribe T7 High Yield RNA Synthesis Kit (NEB) according to the manufacturer’s instructions, and the resulting RNAs were isolated from the reaction mixtures by Monarch RNA Cleanup Kit (NEB). The size and purity of the RNA constructs was confirmed by TBE-urea PAGE (**Supplementary Fig. 3**). Due to the differences in electrophoretic mobility, the sizes of the RNA constructs were additionally confirmed by MALDI-TOF (**Supplementary Fig. 4**). All RNA samples were stored at -80°C before use.

The rPeptides were synthesised by the Spyder Institute using standard protocols for solid-phase peptide synthesis. The identities and purities of the peptides were confirmed by mass spectrometry using UltrafleXtreme MALDI-TOF/TOF mass spectrometer (Bruker Daltonics, Bremen, Germany) according to the standard procedure (**Supplementary Fig. 5**).

### Peptide structural characterization

Circular dichroism (CD) spectra of rPeptides were recorded using a Chirascan-plus spectrophotometer (Applied Photophysics, Leatherhead, UK) over the wavelength range 190– 260 nm in steps of 1 nm with an averaging time of 1 s per step (**Supplementary Fig. 6**). Cleared peptide samples at 0.2 mg/mL concentration in 10 mM Tris (pH 7.5), 1 mM KCl, 1 mM MgCl_2_, 1 mM CaCl_2_ were placed in 1 mm path-length quartz cells, and spectra were recorded at room temperature. The CD signal was obtained as ellipticity in units of millidegrees, and the spectra were averaged from two scans and buffer-spectrum subtracted. The resulting CD spectra were then expressed as molar ellipticity per residue *θ* (deg·cm^2^·dmol^−1^). All CD measurements were performed twice for every rPeptide and curves were then averaged.

### Characterization of rPeptide-rRNA interaction by microscale thermophoresis

In order to determine the dissociation constant of rPeptide-rRNA complexes, the 3’-end of the RNA constructs was fluorescently labelled with fluorescein-5-thiosemicarbazide (FTSC) using a published protocol (17) with slight modifications. First, 500 pmol of PTC rRNA was oxidised by mixing with 10 nmol NaIO_4_ (20 eq.) and 100 mM potassium acetate (pH 5.2) to a total volume of 50 μL and incubating for 90 min at room temperature in the dark. The oxidised PTC rRNA was subsequently purified from the reaction mixture with Monarch RNA Cleanup Kit (NEB) and immediately mixed with 150 nmol FTSC (300 eq.) dissolved in DMF and 100 mM potassium acetate (pH 5.2) to a total volume of 50 μL. The reaction mixture was incubated overnight at 4°C in the dark, and the excess of FTSC was removed with Monarch RNA Cleanup Kit (NEB).

The interaction of the five rPeptides with both WT PTC variants was measured using microscale thermophoresis (MST). Experiments were conducted in duplicate on a Monolith NT.115 system (NanoTemper Technologies). To estimate the effect of metal ions, rPeptide solutions were prepared in 20 mM Tris-HCl (pH 7.5), while PTC rRNA solutions were prepared in two conditions: (i) 20 mM Tris-HCl (pH 7.5) or (ii) 20 mM Tris-HCl (pH 7.5), 2 mM KCl, 2 mM MgCl_2_, 2 mM CaCl_2_. Prior to the experiment, PTC RNAs were heated at 85°C for 30 s and then slowly cooled to room temperature over 30 min, to allow the molecules to refold.

A 2-fold dilution series of the unlabeled rPeptides was prepared with the concentrations of rPeptides ranging from 1 mM to 61.04 nM. 10 μL of rPeptide solutions were subsequently mixed with 10 μL of 100 nM fluorescein-labelled PTC rRNA and the samples were incubated for 15 min at room temperature. Following incubation, the samples were filled into standard-treated capillaries (NanoTemper Technologies) and subjected to MST analysis. The MST measurements were performed at 1 % LED power (green channel) and 40 % MST power, laser-on time 20 s, laser-off time 5 s. The results were analysed by TJump analysis, and the normalised fluorescence values were plotted against the rPeptide concentration. Dissociation constants (**Figure 2**) and binding stoichiometries were then estimated adopting both a single-site model and a cooperative Hill model to fit the curves (**Supplementary Fig. 7**).

**Figure 2:**
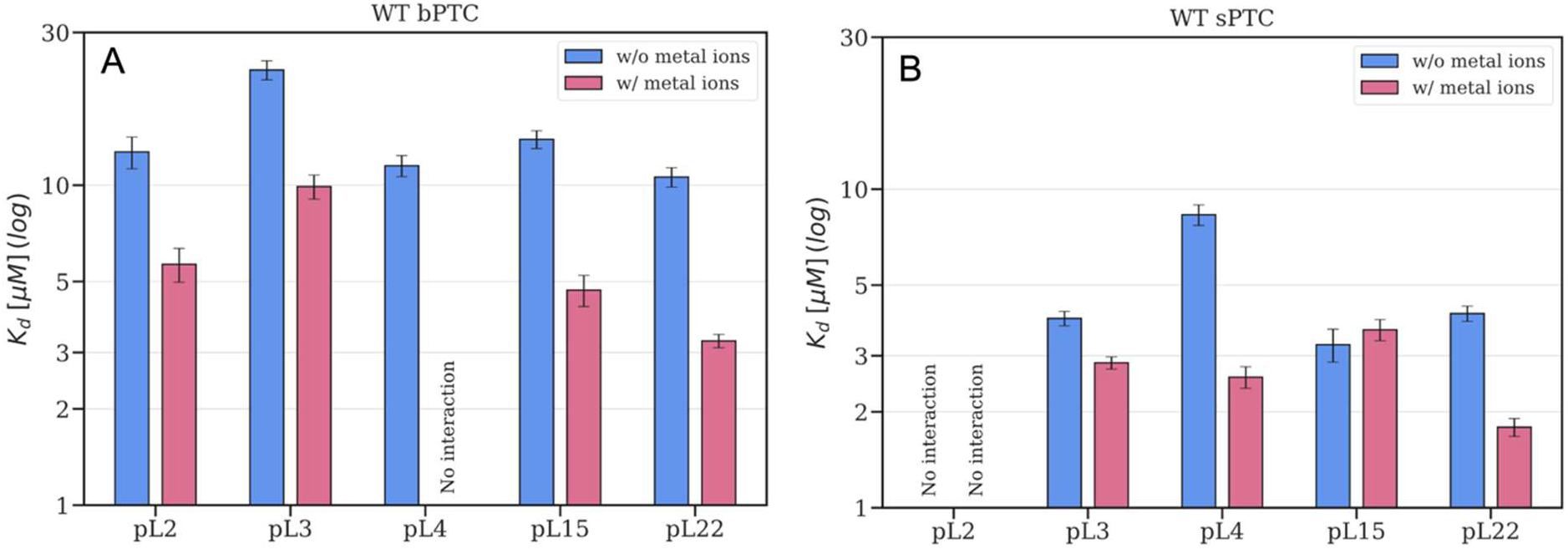
Summary of the WT bPTC (*A*) and WT sPTC (*B*) dissociation constants (*K*_*d*_) with the rPeptides determined using MST in the absence (*blue*) and presence (*red*) of metal ions (1 mM Mg^2+^, K^+^ and Ca^2+^). Error bars represent the semi-dispersion of the two replicates; “no interaction” indicates the cases where the potential *K*_*d*_ value is outside the measurable range of 30 nM to 500 μM or the dissociation curve could not be fitted.

### Molecular dynamics simulations

Several molecular systems were studied using all-atom molecular dynamics (MD) simulations, namely WT bPTC, WT bPTC + rPeptides (pL2, 3, 4, 15 and 22), WT sPTC, WT sPTC + pL2, WT sPTC + pL3, and WT sPTC + pL4. The initial atomic coordinates were obtained by shaving the X-ray structure of *T. Thermophilus* ribosome (PDB 4V51) (18) and interconnecting the flanking rRNA ends by stem loops as detailed in (9) and (11). The sequences of rPeptides were adjusted to match our experiments. Structural Mg^2+^ ions were used as found in the X-ray structure within the cut-off distance of 0.5 nm from the selected regions. The systems were placed in a periodic rhombic dodecahedral box ensuring the distance of 1.5 nm between the box face and the solute. The box was filled with explicit water molecules and K^+^ and Cl^-^ ions of the concentration 150 mM. Finally, each box was neutralised by adding a sufficient amount of K^+^ ions.

The system was described by the Amber family of force fields: ff10 for rRNA (19) (20), ff12SB for peptides (21), Joung and Cheatham ions (22), and SPC/E water (23). The systems were energy minimised in about 80000 steps of the steepest descent algorithm and subsequently heated to 300 K during 100 ps simulations. The initial velocities were randomly drawn from the Maxwell-Boltzmann distribution at 10 K. The pressure and density were equilibrated during another 100 ps simulations. In the course of minimization and equilibration, *position* restraints (*k* = 1000 kJ mol^−1^ nm^−2^) were applied to all heavy atoms of the solute to prevent its spurious conformational changes. The unrestrained production runs were carried out at constant temperature of 300 K and pressure 1 bar, using a v-rescale thermostat (24) and Parrinello-Rahman barostat (25). The long-range electrostatics were treated by particle mesh Ewald algorithm; van der Waals interactions were described by the Lennard-Jones potential with the cut-off of 1.0 nm. A virtual-site algorithm was used to hydrogen atoms allowing for the integration time step of 4 fs. Three independent trajectories were obtained for each system differing in the initial set of velocities. We collected trajectories of 1100 ns each with trajectory frames saved every 100 ps.

Prior further analyses, all trajectories were superimposed with respect to the largest common rRNA fragment (113 nt, **Figure 3D**) making the bPTC and sPTC constructs better comparable. A structural analysis involved the last 300 ns of each trajectory. The atomic coordinates were averaged over the simulation time and the three independent trajectories, which provided MD-based structural models of the WT constructs. Structural differences of the models were represented by Euclidean distances between equivalent atoms of the MD models. For the fluctuation analysis, the first 300 ns of each trajectory were skipped and the root-mean-square fluctuations were calculated over the rest of the simulation time. The RMSFs were averaged over the independent trajectories and projected onto the initial structure (**Figure 3B-C**).

**Figure 3:**
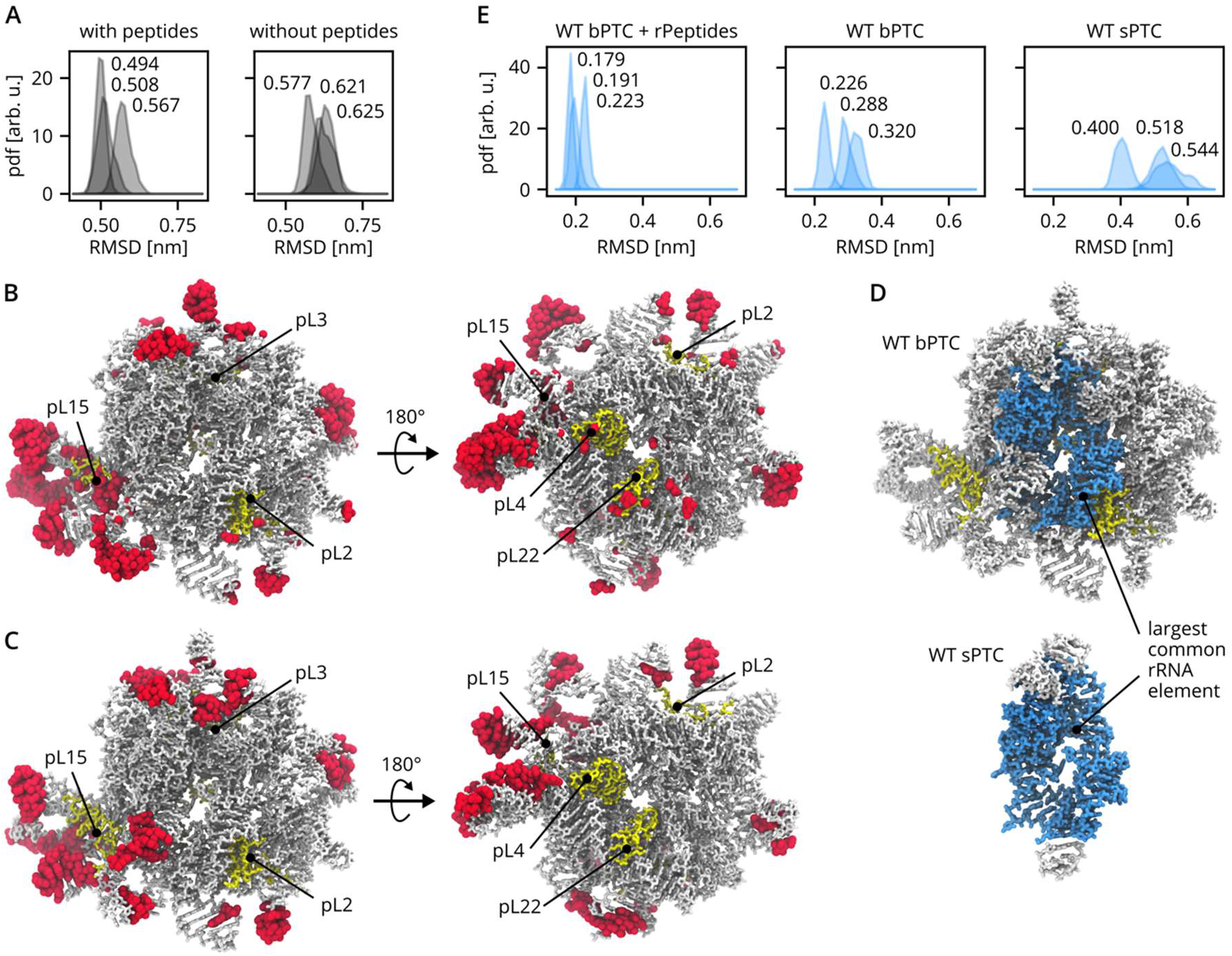
Overview of MD simulations. (A) Probability density functions (pdfs) of root-mean-square deviations (RMSD) calculated for WT bPTC phosphorus atoms with respect to the initial conformation. The last 300 ns of each independent trajectory were used. Mean values of pdfs are denoted. (B) The most flexible residues as identified by root-mean-square fluctuation (RMSF) analysis of the last 800 ns of each trajectory. Top 10% residues with the highest RMSF are shown in red, rRNA in white, rPeptides in yellow. The two viewpoints show the tRNA binding site and the ancestral exit tunnel lined by pL4 and pL22. For clarity, the location of pL3 is shown, although it is buried in the rRNA. (C) Same as (B), but showing the rRNA residues identified as the most structurally different between MD models obtained with and without rPeptides. Top 10% residues with the largest structural difference are in red. (D) The largest common rRNA element of WT bPTC and sPTC in blue. (E) Pdfs of RMSD of the largest common rRNA element. The last 300 ns of each independent trajectory were used. Mean values of pdfs are denoted.

### Coacervation

RNA solutions were prepared in RNase free water in small aliquots (1 - 20 μL) and at various concentrations, and they were stored at -80°C. Lyophilized rPeptides were suspended in RNase free water to typical concentrations of 50 mM. These stock solutions were further diluted at different concentrations, divided in small aliquots and stored at -20°C. Phase separation diagrams were estimated as a function of the *R*_+/-_ ratio and of the molar charge concentration (in units of the elementary charge) *Q* of the mixtures (**Figure 4** and **Supplementary Fig. 8 & 9**), defined as:

**Figure 4:**
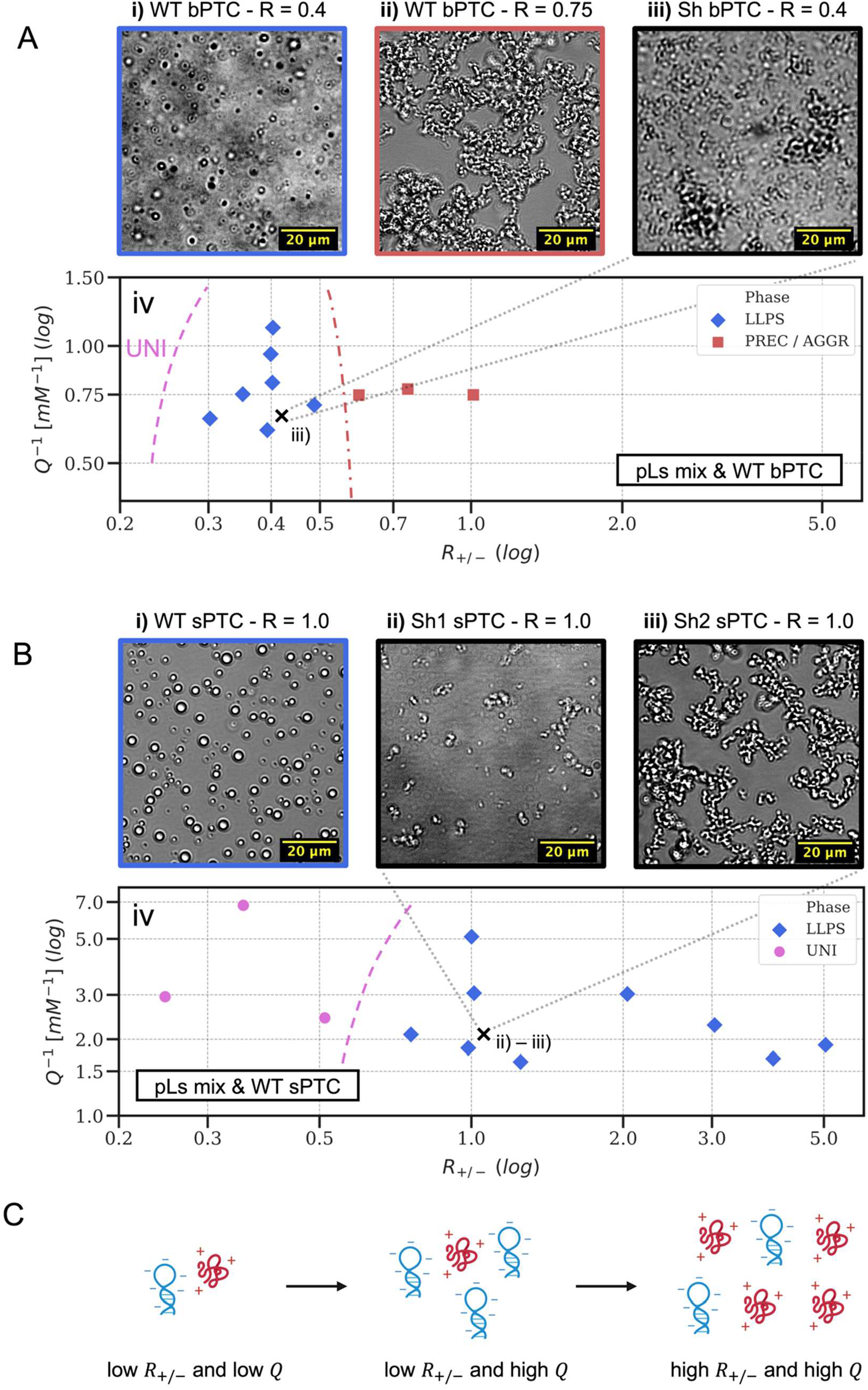
Phase separation of WT bPTC (*sector A*) and WT sPTC (*sector B*) with equimolar mixtures of pL2, 3, 4, 15 and 22, compared to the phase behaviour of the shuffled variants Sh bPTC, Sh1 sPTC and Sh2 sPTC (*panel iii* in *sector A* and *panels ii & iii* in *sector B*, respectively, see **Supplementary Fig. 2**). Phase diagrams (*panels iv* in *sector A & B*) are drawn as a function of the stoichiometric charge ratio *R*_+/-_ (x axis, see *eq. 1* in **Mat. & Meth**., ***coacervation***) and of the inverse of the molar concentration of charged units *Q*^−1^ (y axis, see *eq. 2* in ***ibidem***), with both axes in log scale. Different colours and markers are associated to different phases, with “UNI” indicating a uniform solution, “PREC/AGGR” the presence of precipitates or aggregates and “LLPS” the formation of liquid-like droplets. Dashed curved lines in the phase diagrams are arbitrary guidelines for the eye and mark the region where no demixing (“UNI”) or precipitation/aggregation is observed. The series of optical micrographs in *sector A* (*panels i, ii, iii*) shows 2 distinct phases for the WT bPTC with colours matching the phase legend and compares the phase behaviour of the WT bPTC at *R*_+/-_ = 0.4 (*panel i*) to the one of the Sh bPTC at the same *R*_+/-_ value (*panel iii*). The “X” cross on the diagram marks the spot related to the Sh bPTC measure. The series of optical micrographs *in sector B* (*panels i, ii, iii*) compares the LLPS of the WT sPTC at *R*_+/-_ = 1 (*panel i*) to the phase behaviour of Sh1 and Sh2 sPTC (*panels ii & iii*, respectively) at the same *R*_+/-_ value. The “X” cross on the diagram marks the spot related to the Sh1 and Sh2 bPTC measurements. *Sector C* shows a pictorial representation of the *R*_+/-_ and *Q*^−1^ parameters for a mixture of a single peptide (*in red, carrying +3 charges*) and an RNA molecule (*in blue, carrying -6 charges*).

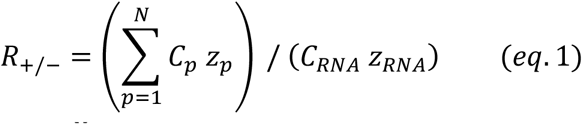

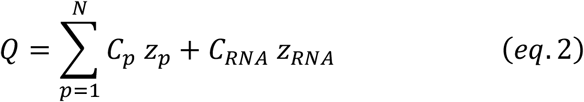

where *C*_*p*_ and *C*_*RNA*_ are the molar concentrations of rPeptides and rRNA, respectively, and the summation runs over the different rPeptides employed. The *R*_+/-_ ratio represents the stoichiometric charge ratio of the molecules in solution (positive over negative charges), while *Q* denotes the molar concentration of charged units. *Z*_*RNA*_ is defined as the length of the strand minus 1 (the RNA terminals are not phosphorylated), while the net peptide charges (*Z*_*p*_) are estimated using the pH 7.5 values given by the *Peptide Property Calculator* by NovoPro (https://www.novoprolabs.com/tools/calc_peptide_property): pL2 = +4.1, pL3 = +5.0, pL4 = +4.0, pL15 = +5.0, pL22 = +7.0. The typical *R*_+/-_ values explored in the phase separation diagrams range from 0.1 to 5, while *Q* ranged between 0.1 mM and 2 mM. When performing experiments in which more than one rPeptide was involved, all the peptide concentrations were set equal and the desired *R*_+/-_ and *Q* values were tuned by changing the RNA and the overall common peptide concentration.

In order to explore different values of *R*_+/-_ and *Q*, mixtures of the WT bPTC and WT sPTC and of pL2, 3, 4, 15 and 22 were prepared at various working concentrations, at room temperature and by keeping the ionic strength fixed. The same buffer was used in all the experiments: 20 mM Tris pH 7.5, 1 mM KCl, 1 mM MgCl_2_, 1 mM CaCl_2_. The ionic strength of the buffer, accounting both for the Tris ionic strength and for the dissociated monovalent K^+^ and Cl^−^ and divalent Mg^2+^ and Ca^2+^ ions, is estimated to be 23 mM in equivalent NaCl concentration. Mixtures of pL22 with either WT bPTC and WT sPTC were prepared by adding, in this given order, 1 μL of 8x buffer, 3.5 μL of peptide and 3.5 μL of RNA, while mixtures of more than one rPeptides with RNA were prepared in total volumes of 8 μL by adding the same amount of each peptide, adjusting with 1 μL of 8x buffer and by final addition of RNA. All the necessary dilutions were carried out in RNase free water and samples were prepared in small PCR plastic tubes. Before adding it to the peptides-buffer mixture, the RNA was annealed in a thermal cycler from 85°C to 40°C in 30 minutes in 3°C steps to refold the molecules. The RNA solution was then left reaching room temperature before mixing it to the rPeptide(s).

Liquid-liquid phase separation (LLPS) phase diagrams were characterised through bright field microscopy. For imaging, samples were transferred in a plastic multiwell which was previously treated with a BSA 3% in mass solution (the BSA solution was added to each well, removed after 20 min and then washed two times with water). Well plates were sealed with PCR plastic films and microscopy measurements were carried out at room temperature. Samples were inspected at the Imaging Methods Core Facility at BIOCEV using a Leica DMi8 WF microscope. Images were taken at different magnifications (10x, 20x, 40x) and the phase state was annotated for further characterization of the phase diagrams.

### Testing of rRNA stability

A solution of 2 μM WT sPTC RNA was incubated with or without pL22 (39.0 μM, R=1.0) or pL22 = (10.5 μM, R=0.27), or a mixture of pL2, 3, 4, 15, and 22 (10.5 μM of each peptide, R=1.0) in Buffer A (20 mM Tris-HCl pH 7.5, 1 mM KCl, 1 mM MgCl_2_, 1 mM CaCl_2_) in the total volume of 10 μL. The 2 μM solution of WT bPTC was incubated with or without pL22 (70.3 μM, R=0.4) or pL22 = (18.8 μM, R=0.1), or a mixture of pL2, 3, 4, 15, 22 (18.8 μM of each peptide, R=0.4) in Buffer A. In order to perform the RNase stability test, mixtures were divided into halves and incubated for 1 h at room temperature. Monarch RNase A (NEB) (0.1 ng/mL in final concentration) was added, and both samples with and without RNase A mixtures were incubated at 37°C for 30 or 40 min.

The reactions were stopped by adding 80 units/μL of “Proteinase K, Molecular Biology Grade” (NEB) and incubated for 5 min at 3°C. Reaction products were analysed using 10% Urea PAGE. Gels were stained by GelRed (Biotium) and visualised by UV. Band intensities were evaluated by Fiji (Schindelin). The experiments were performed in triplicates and statistical analyses were done using Excel 2023 (Microsoft). Comparisons between datasets were performed using a two-tailed Wilcoxon-Mann-Whitney test with n = 3. A p-value of *≦* 0.2 and u = 0 was considered to be a statistically significant difference.

## Results

### rPeptides are devoid of structural elements and interact with PTC rRNA

The WT bPTC directly interacts with a number of rPeptides within the ribosome and their parts are intertwined within the RNA structure (**Figure 1A**). The WT sPTC, on the other hand, is much less dependent on these rProtein fragments (**Figure 1A vs. 1B**). To resolve their role during ribosomal evolution, the peptide sequences resembling these rPeptides were synthesised and purified (**Supplementary Fig. 5**). Some of the sequences (pL2, pL3 and pL22) were adapted precisely from a previous study (9), while pL4 and pL15 were truncated based on our structural analysis to best preserve their bPTC binding properties and to allow for the solid-phase synthesis of all of them (**Figure 1C**). All the rPeptides are well soluble and their CD spectra confirm a general lack of secondary structure arrangement (**Supplementary Fig. 6**).

The interaction of the five rPeptides with both WT PTC variants was measured using MST. The 3’ end of the PTC rRNA was labelled by fluorescein-5-thiosemicarbazide and purified. To refold its three-dimensional structure, PTCs were kept heated at 85°C for 30 seconds and gradually cooled to room temperature. The refolded PTCs were mixed with different concentrations of the rPeptides to detect dissociation constants (*K*_*d*_). This was done in both the absence and presence of 1 mM Mg^2+^, K^+^ and Ca^2+^; the choice of these ions is based upon their prebiotic abundance and prevalence in extant ribosomal structures. We hypothesised that they may play an important role in tuning the interaction between heavily charged molecules, such as Arg-rich peptides and RNA.

Both PTC variants interact with most of the rPeptides with *K*_*d*_ in μM range (**Figure 2** and **Supplementary Fig. 7**). Dissociation constants could not be estimated only for pL2 with WT sPTC and for pL4 with WT bPTC in presence of metal ions. While the binding curves remain almost unchanged for the WT sPTC interaction with rPeptides upon addition of metal ions, the WT bPTC interaction with most of the rPeptides is visibly affected. Based on the shape of the binding curves and Hill coefficient estimates, rPeptides (with the exception of pL4) seem to bind to WT bPTC with a decreased peptide:RNA stoichiometry in the presence of metal ions, with *K*_*d*_ values decreasing 2-3 times (**Figure 2** and **Supplementary Fig. 7**).

### rPeptides stabilise the bPTC fold while the sPTC is more flexible and structurally different from the bPTC

The structural properties of the PTC constructs and the effects of the rPeptides were further explored by all-atom MD simulations. First, we performed a series of MD simulations of WT constructs to check their conformational flexibility and interactions of their components. The simulations were started from the three-dimensional structures as found in the *T. Thermophilus* ribosome, so our simulations report on the (dis)similarity of the constructs to the modern ribosome.

In the course of microsecond-long simulations of WT bPTC with rPeptides, all of the rPeptides remained well associated with the rRNA, suggesting low dissociation rates. We observed no large-scale conformational changes of the rRNA, as illustrated by the root-mean-square deviation (RMSD) with respect to the initial conformation (**Figure 3A**). Fluctuation analysis showed that the most variable regions (highlighted in red) are the stem loops, peptide pL15 and some residues in the ancestral ribosome tunnel (**Figure 3B**). In addition, we performed MD simulations of WT bPTC without all rPeptides. A comparison of the two bPTC systems revealed that the rPeptides reduce RMSD of the rRNA, thus making it conformationally more similar to the modern ribosome (**Figure 3A**). However, the largest structural differences of rRNA related to the presence/absence of the rPeptides in the WT bPTC are located mostly on the surface of the construct (**Figure 3C**).

Compared to WT bPTC, the conformational freedom of WT sPTC was much broader as described by RMSD of the largest common rRNA element present in both bPTC and sPTC (113 nt, **Figure 3D & E**). Also, the RMSD reached higher values in WT sPTC, so the structure was less similar to the modern ribosome than the WT bPTC. Overall, the simulations indicate that the WT constructs may adopt compact tertiary structures similar to the modern ribosome and that the tertiary structures depend on the presence/absence of rPeptides and the size of the rRNA.

### rPeptides drive coacervation of protoribosome constructs

While WT bPTC and WT sPTC are fully soluble in the buffer when in solution as single species, the addition of equimolar mixtures of rPeptides (verified to be soluble at the same average concentrations used in rPeptides-RNA mixtures) triggers a rich phase behaviour, including precipitation of solid particles, formation of branched aggregates and demixing into liquid-like droplets (**Figure 4, *micrographs A i-ii & B i***). In particular, we investigated the onset of LLPS, or coacervation, as a function of the total concentration of RNA and peptides on the y axis (expressed in **Figure 4A iv & 4B iv** as the inverse of the molar charge concentration) and of their ratio (peptides over RNA) on the x axis, namely *R*_+/-_ (expressed as the ratio of positive and negative charges, see **Mat. & Meth**., ***coacervation***, *eqs. 1 & 2*), as sketched in **Figure 4C**. For both WT bPTC (**Figure 4A iv**) and WT sPTC (**Figure 4B iv**), coacervation is observed at moderate total concentrations (as low as 2 μM RNA) and at high enough relative concentration of rPeptides (*R*_+/-_ > 0.3 and 0.7, respectively). However, bPTC coacervation is suppressed for *R*_+/-_ > 0.75, while it is observed for sPTC over a wide range of *R*_+/-_ values. Qualitatively similar results are obtained for mixtures of both protoribosomes with a single rPeptide, namely pL22 (the rPeptide with the highest ratio of positively charged residues), corroborating the idea that the phase behaviour is determined by the different rRNA constructs involved (**Supplementary Fig. 8**). In addition, we observed that WT sPTC coacervates with pL22 are stable up to 90°C: this is consistent with an “upper critical solution” phase behaviour, where enthalpic interactions are the main drivers of LLPS.

### The coacervation of rRNA-rPeptide protoribosomes is dependent on rRNA structure and composition

The complexation of positively charged rPeptides and negatively charged PTC is clearly at the basis of the observed LLPS, as previously reported for a variety of other nucleic acids-amino acids systems (26) (27) (28) (29). However, we found that the sequence composition of the employed RNA strands, as well as their secondary and tertiary structures, sensitively affect coacervation propensity. We mixed rPeptides with two sPTC variants of shuffled sequences, so that the predicted fraction of paired bases was either higher or lower than in the original WT sPTC and the symmetric structure of the sPTC was disrupted (namely Sh1 sPTC and Sh2 sPTC, respectively: **Supplementary Fig. 2**). Similarly, a single shuffled variant (Sh bPTC) was designed also by permutation of WT bPTC sequence. In such cases, substantial weakening of the LLPS propensity was found, as shown by the optical *micrographs A iii, B ii-iii* compared to *A i* and *B i*, respectively, in **Figure 4**, corresponding to the inner LLPS regions for WT bPTC and sPTC, marked by “X”. The dissimilarity in the phase behaviour extends well beyond these points, as shown in **Supplementary Fig. 9** for the Sh bPTC variant: aggregation or precipitation occur at all examined samples in the (*R*_+/-_, *Q*^−1^) space, highlighting the subtle dependence of LLPS on RNA secondary and tertiary structure.

To further examine the effect of nucleotide composition, various unstructured single-stranded DNA oligomers with different A-T-C-G compositions were mixed with the pL3 peptide. The utmost LLPS propensity occurs when the nucleotide composition closely resembles the native rRNA nucleotide environment of the pL3 peptide (namely: A = 10.7%, C = 39.3%, G = 32.1% U = 17.9%) and LLPS stability monotonically decreases as the composition deviates from it, as shown in **Supplementary Fig. 10**. This is intriguing, as the rRNA environment sequence composition is different for each rPeptide and generally not very rich in purine bases, which have been previously shown to favour LLPS (29).

### The coacervation effectively increases the stability of the PTC rRNA

Inside the coacervate, PTC and peptides are condensed and separated from a dilute aqueous solution, suggesting that coacervation may help to stabilise or protect RNA from hydrolysis. To investigate this, the PTC stability was tested against enzymatic degradation.

RNase A was added to the WT bPTC and sPTC with or without rPeptides in concentration ratios corresponding to the inner LLPS region in the phase diagrams in **Figure 4**. The rPeptides were either used in combination or the same R ratio (defined by *eq. 1* in **Mat. & Meth**., ***coacervation***) was obtained only using pL22 (previously used for testing LLPS propensity, see **Supplementary Fig. 8**). The potential degradation of PTCs under these conditions was visualised by Urea-PAGE (**Figure 5A & B**, *lanes 1-6*). While the bPTC construct was not significantly protected by the rPeptides against enzymatic degradation under any of the tested conditions, a robust sPTC stabilisation and protection was found when either pL22 alone or rPeptide mixture was used in the condition.

**Figure 5:**
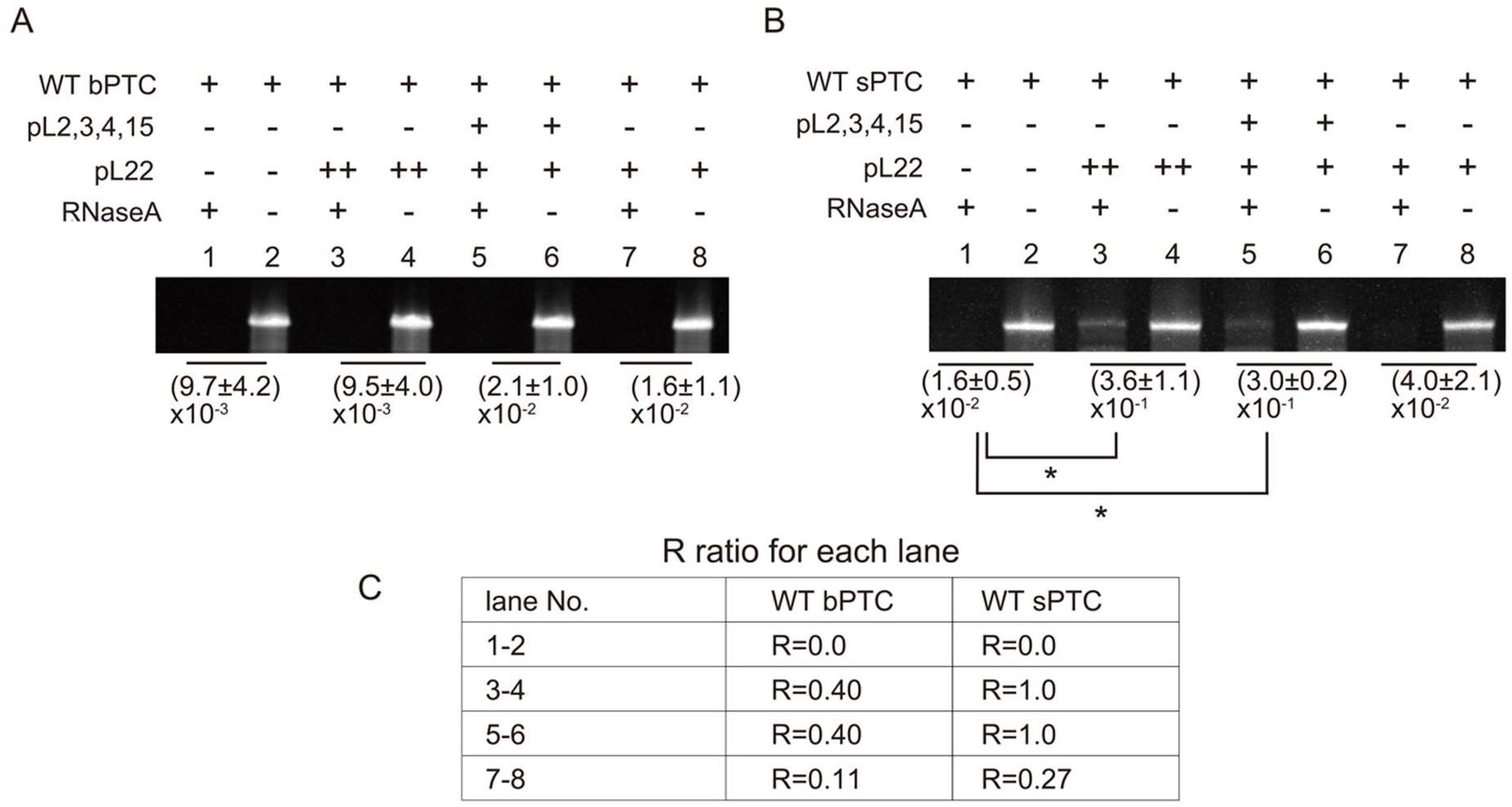
rRNA stability to RNase A treatment in the absence and presence of rPeptides. (A) WT bPTC and (B) WT sPTC. The average ratios and standard deviations of the enzymatically treated vs. untreated RNAs are listed below the respective bands, along with their significance values where appropriate. (* = p-value less than 0.2, n = 3 and u = 0 Wilcoxon-Mann-Whitney test). (C) Table of R ratios in each lane.

To test the importance of the coacervation-prone molar ratios, the pL22 peptide was used also in a significantly smaller concentration (∼4x), representing its molar contribution from the rPeptide combination (**Figure 5A & B**, *lanes 7-8)*. The sPTC was fully degraded under such conditions, supporting our hypothesis that coacervation, and not just interaction, is needed for the protection against enzymatic degradation.

## Discussion

Following the past decades of the RNA-first (or other “Me-First”) narratives in the origins-of-life, discussions have gradually shifted towards a greater emphasis on the systems chemistry level of thinking (13) (30) (31). Several recent studies have succeeded in syntheses of molecules and proto-biological structures that could be nominated for possible routes towards early life (32) (33) (34) (35) (36) (37). Nevertheless, to truly understand the origin of *our* life (vs. possible routes to life), gaps must be filled between the prebiotic chemistry, protobiology and ancestral life forms. The PTC (or protoribosome) clearly represents one of the most profound molecular fossils of ancestral life. It is one of the farthest points that we can reach by tracing back current molecular life. Understanding how it came about and its route towards early biology therefore provides a unique opportunity to fill these gaps.

Two groups have recently reported the PTC activity using RNA structures reconstructed from the core of extant ribosomes (10) (11) (12). The RNA itself was therefore shown to be capable of peptide bond formation between amino acids, albeit with very low efficiency. Nevertheless, due to the prebiotic abundance of amino acids and the facile nature of their condensation (e.g. by wet-dry cycling), short peptide sequences were most probably prebiotically plausible also in the absence of ribosomal synthesis (13) (38). The role of peptides in the emergence and evolution of the protoribosome is the subject of this study.

We reproduced two previous models of the PTC rRNA constructs: (i) the sPTC resembling a more compact and ancestral form that has been shown to possess the peptidyl-transferase activity, and (ii) bPTC representing a ∼4.5x larger construct that – within the natural ribosome – interacts with the fragments of several ribosomal proteins. Using MD simulations, we observe that the sPTC displays much broader conformational freedom, while the bPTC is more rigid and structurally similar to the modern ribosome. Our observation of the bPTC structural properties broadly agrees with its previous characterization (9).

To examine the role of peptides in the different proto-ribosomal scenarios, we further included five rPeptides (fragments of proteins that interact with the PTC in native ribosome) in our study. These peptides contain all canonical amino acids and are enriched in positively charged residues. Diverse lines of evidence suggest that prebiotically available amino acids did not include about half of the extant alphabet, missing e.g. today’s positively charged and larger functional groups, such as aromatics (39) (40) (41). The five rPeptides therefore do not represent the most prebiotically plausible peptide variants; they can rather be viewed as peptide fossils that have been preserved in the ribosomal core. All five rPeptides are devoid of regular secondary structure arrangement.

We observe that most of the rPeptides interact with both rRNA PTC constructs with affinities in μM range. In case of bPTC, the binding curves change and dissociation constants decrease 2-3 times upon addition of metal ions which are (i) found in extant ribosomal structures, (ii) considered prebiotically plausible, and (iii) have been previously shown to stabilize the PTC (42). In contrast, the rPeptide-sPTC binding curves are almost unaffected by the metal ion addition. The presence of metal ions may play an important role in minimising non-specific binding which would be based on the significantly opposite charged components of the interaction partners only. The inclusion of the rPeptides in the MD simulation of the bPTC construct further reduced the RMSD and suggested that the rPeptides could have played a role in stabilisation of the rRNA structure in the protoribosome. MD simulations also suggested that the interactions of the rPeptides with the sPTC are less specific than with the bPTC. Although we did not observe the dissociation of the rPeptides from the sPTC within the simulation time, pL2, pL3, and pL4 were all two- to three-fold more mobile on the sPTC than on bPTC, as characterised by the RMSD and conformational ensembles of the rPeptides (**Supplementary Fig. 12**).

Similar (although in some cases longer) rPeptide sequences were used previously by Hsiao et al. to characterise the properties of the bPTC construct (9). They observed binding of pL4, pL3, pL15 and pL22, where only pL4 binding was tested in the peptide form and the remaining peptides were prepared as fusions with the maltose-binding protein. Intriguingly, in that study all the binding tests were performed in the absence of metal ions, i.e. conditions which implied less specific interaction properties in our study (9). In our experimental setup, the pL4 peptide (shorter by 15 amino acids than in the previous study) is the only one that does not display any interaction upon metal ion addition.

Finally, the addition of rPeptides to the PTC rRNA constructs was observed to induce LLPS of both construct variants. However, the bPTC liquid demixing is limited to a narrow area of the phase diagram, while the sPTC construct displays LLPS under a broader range of peptide/rRNA concentrations and is qualitatively more prominent. We hypothesised that the PTC coacervation may have a stabilising effect on the rRNA moiety and confirmed that the sPTC rRNA construct was protected against enzymatic degradation. This was true when the experiments followed the LLPS-prone concentration ratios, and the effect was therefore LLPS-based rather than interaction specific. The LLPS propensity of the PTC constructs was also observed to be rather sensitive to the structure and sequence of the rRNA.

The protoribosome (sPTC) was previously shown to possess the peptidyl-transferase activity (11). Here, we show that the arguably most ancient fragments of the ribosomal proteins trigger the protoribosome coacervation under a wide range of concentrations, providing compartmentalization and stabilisation of the sPTC construct against degradation. Our results demonstrate a clear example of a “peptide-RNA” co-evolution using the very fossil of our biological life as an example. Several studies have recently evoked Oparin’s century-old theory that coacervate microdroplets could have played major roles during life’s origins (16) (26) (43). Because of the high RNA-peptide LLPS propensity, coacervates have been proposed as a unique environment supporting co-evolution of those species (16). Current studies are now researching the impact of different environments as well as the polymer properties and lengths on the coacervation propensities (43) (44). The bigger construct used in our study, bPTC, is structurally more rigid and interacts with the rPeptides with higher specificity. Compared with sPTC, it has only weak LLPS propensity and the LLPS conditions do not provide additional stability to bPTC.

Based on the data presented here, we speculate that during origins of life, coacervation could be more prominent with flexible structures and based on interactions that could have lacked specificity. Such compartmentalization could lead to concentration of reactants and enhance the activity as previously demonstrated using different ribozyme examples (45) (46). It could also lead to evolution of ribosomal peptides towards sequences that could interact with the gradually accrued rRNA structures more specifically and to help in such structure stabilisation and sovereignty. Nevertheless, to provide the true link between prebiotic chemistry and the biological past, it will be of key importance to test these hypotheses using prebiotically plausible pre-ribosomal peptides as well as compare the potential impact of pre-/ribosomal peptides on the PTC activity.

## Data availability

The data underlying this article are available in the article and in its online supplementary material.

## Supplementary data

Supplementary Data are available at NAR online.

## Acknowledgements

We thank Ing. Dalibor Pánek for his technical support at the IMCF at BIOCEV campus and Dr. Vyacheslav Tretyachenko for helpful discussions about this work.

