## Supplementary material for "The interplay between peptides and RNA is critical for protoribosome compartmentalization and stability"

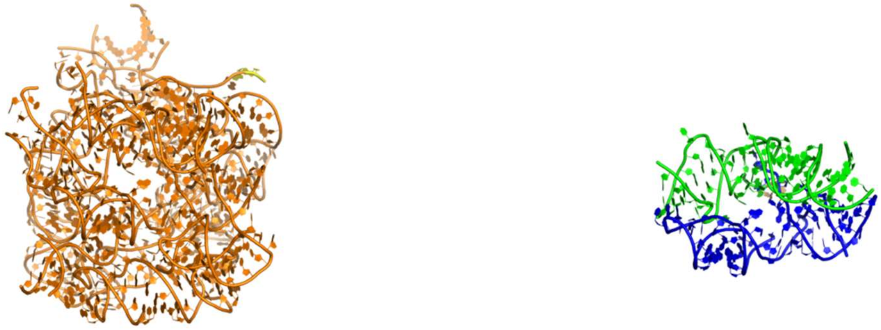

big PTC

```
1        10        20        30        40        50
GGGUGCCUAU UGAAGCAUGA GCCGGCGACU CAGCCGUAAG GCUGGACCCG
AAACCGGGCG AGCUAGCCCU GGCCAGCCGU AAGGCGAACC GGUGGGGGAU
GCAAACCCCU CGGAUGAGCU GGGGCUAGGA GUGAAAAGCU AACCGAGCCC
GGAGAUAGCU GGUUCUGCCG UAAGGCAGCG UUGCCGUAAG GCAAGUGCGA
AUGCCGGCAU GAGUAACGAG CGGGAGAACC CUCGCCAAGG AACUCUGCAA
GCCGUAAGGC CUCUGGCGAC UGUUUACCAA AAACACAGCG CCGUAAGGCG
CGAGCCGUAA GGCUAACGAC CGGAGCGCUG UCUCGGCGAG GGACCCGGUG
AAAUUGAACU GGCCGUGAAG AUGCGGCCUA CCCGUGGCAG GACGAAAAGA
CCCCGUGGAG CUUUACUGCC GUAAGGCAGU UUGACUGGGG CGGUCGGCCG
UAAGGCUAAA AGUUACCCCG GGGAUAACAG GCUGAUCGCC GUAAGGCGGU
UUGGCACCUC GAUGUCGGCU CGUCGCGCCG UAAGGCUUGG GCUGUUCGCC
CAUUAAAGCG GCACGCGAGC UGGGUUCAGA ACGUCGUGAG ACAGUUCGGU
CUCUAUCCGC CACGGGCA
                   617
```

small PTC

```
1        10        20        30        40        50
GGAAGACCCC GUGGAGCUCU UCGGAGUUAC CCCGGGGAUA ACAGGCUGAU
CUCUUCGGAG GUUUGGCACC UCGAUGUCGG CUCGUCGCUU CGGCGGCACG
CGAGCUGGGU UCAGAACGUC GUGAGACAGU UCGGUC
                                      136
```

**Supplementary Fig. 1**. Tertiary structures derived from PDB 4V51 and related sequences of the two WT PTC constructs employed in this study: the 617 nt bPTC (*left panel*) and the 136 nt sPTC (*right panel*). WT bPTC is made up of the 505 nt from the 23S rRNA (*orange*), joined by 11 stem loops (*grey*). The WT sPTC is composed of 116 nt from the 23S rRNA (A region in *green* & P region in *blue*) and connected with 3 stem loops (*grey*). The extra GG for transcription are marked in *yellow*.

**Supplementary Table 1**. DNA oligos used for preparation of PTC RNA constructs.

| Name | Sequence | Note | Reference |
|---|---|---|---|
| WT_bPTC | AAAAAGTGGCCTGTAATACGACTCACTATAGGGTGCC<br>TATTGAAGCATGAGCCGGCGACTCAGCCGTAAGGCTG<br>GACCCGAAACCGGGCGAGCTAGCCCTGGCCAGCCGTA<br>AGGCGAACCGGTGGGGGATGCAAACCCCTCGGATGAG<br>CTGGGGCTAGGAGTGAAAAGCTAACCGAGCCCGGAGA<br>TAGCTGGTTCTGCCGTAAGGCAGCGTTGCCGTAAGGC<br>AAGTGCGAATGCCGGCATGAGTAACGAGCGGGAGAAC<br>CCTCGCCAAGGAACTCTGCAAGCCGTAAGGCCTCTGG<br>CGACTGTTTACCAAAAACACAGCGCCGTAAGGCGCGA<br>GCCGTAAGGCTAACGACCGGAGCGCTGTCTCGGCGAG<br>GGACCCGGTGAAATTGAACTGGCCGTGAAGATGCGGC<br>CTACCCGTGGCAGGACGAAAAGACCCCGTGGAGCTTT<br>ACTGCCGTAAGGCAGTTTGACTGGGGCGGTCGGCCGT<br>AAGGCTAAAAGTTACCCCGGGGATAACAGGCTGATCG<br>CCGTAAGGCGGTTTGGCACCTCGATGTCGGCTCGTCG<br>CGCCGTAAGGCTTGGGCTGTTCGCCCATTAAAGCGGC<br>ACGCGAGCTGGGTTCAGAACGTCGTGAGACAGTTCGG<br>TCTCTATCCGCCACGGGCA | Template of WT bPTC | (1) |
| Sh_bPTC | AAAAAGTGGCCTGTAATACGACTCACTATAGGCCTCG<br>GCGGACCCACCCCAACCTCGAGTAGGGGACCTACACA<br>GAGCATTATCAAACGACAGCCTTCTCAGGTGCATTAA<br>GGACACCAAAAGATCCAGCCGCCGGATGGGATGAGTG<br>ACCGACCGCTACTAGGCCGCATGAGACCCCAATATCC<br>CGTGCAGACAGGGAAAGGCACAGACGAGGTCAGTCAA<br>ACTGGTCCGCTTCACAGGGGGAGTCACTGCAAGGTCG<br>TATACTGCGTGGTTGGGTACCGGTCGGGTCTGCTGAC<br>CAGTATGAGGTATTCCCTACTCTAGGGATGACCGAAG<br>GGTGTTAGTTCCCGAGGAGATTCCCGCCCAGGAATGC<br>TAGATGGACGCGGGAGCGTGTAGTCAGGGGTAAGACT<br>TAGACACGGGTCCACGTAGCGGTGGCACTGATAGGTG<br>CGTTATGCGCCGCGACGCTGGGCTGGGCCAGGTGACG<br>TCGAGCAGTGCTGACGGCGAAGGGCGGCGCCGCGGGC<br>GAGGATCTAATCAGGCTTTACTGTGCAGTAGGACGGG<br>GATGGCCCGGGCACGCTATACGGCGACGCCCGCCCAC<br>GGTAAGCGGGAGAGATGCGACCATCGAGCAGCCGAGT<br>GGTCTATCCGCCACGGGCA | Template of Sh bPTC | This study |

| | | | |
|---|---|---|---|
| bPTC_F | AAAAAGTGGCCTGTAATACGACTCA | Forward primer for preparing dsDNA of bPTC by using DNA polymerase | This study |
| bPTC_R | TCTATCCGCCACGGGCA | Reverse primer for preparing dsDNA of bPTC by using DNA polymerase | This study |
| WT_sPTC_F | GCGTAATACGACTCACTATAGGAAGACCCCGTGGAGCTCTTCGGAGTTACCCCGGGGATAACAGGCTGATCTCTTCGGAGGTTTGGCAC | Forward primer for preparing dsDNA of WT sPTC by using Klenow enzyme | (2) |
| WT_sPTC_R | GACCGAACTGTCTCACGACGTTCTGAACCCAGCTCGCGTGCCGCCGAAGCGACGAGCCGACATCGAGGTGCCAAACCTCCGAAGAG | Reverse primer for preparing dsDNA of WT sPTC by using Klenow enzyme | (2) |
| Sh1_sPTC_R | GCAACCATATGACCAGCCACAACGTCCCATGGACGTTGCATTCACATTGGGCCGTGCCACGGGGCCCTACGGGTCCCGATCCGAGAAATGGCCCCGTGGCTATCGACCATAACTCGGTGCCCCGGAGCAGGCCAATCCTATAGTGAGTCGTATTACGC | Template of Sh1 sPTC | This study |
| Sh2_sPTC_R | CAGACACGCCTGGACGACGACCATGCCAAGTCCCTTACAAGCTTTCAGGCACCCACTACACGCGGGCAATACTGCGATGGCCGCGCTCCTTGTCCGCGGCGACGCACGACACAAGTTCAGGTCTGTTGGGGACCAGCCTATAGTGAGTCGTATTACGC | Template of Sh2 sPTC | This study |
| T7_F | GCGTAATACGACTCACTATAGG | Forward primer for Sh sPTCs | This study |

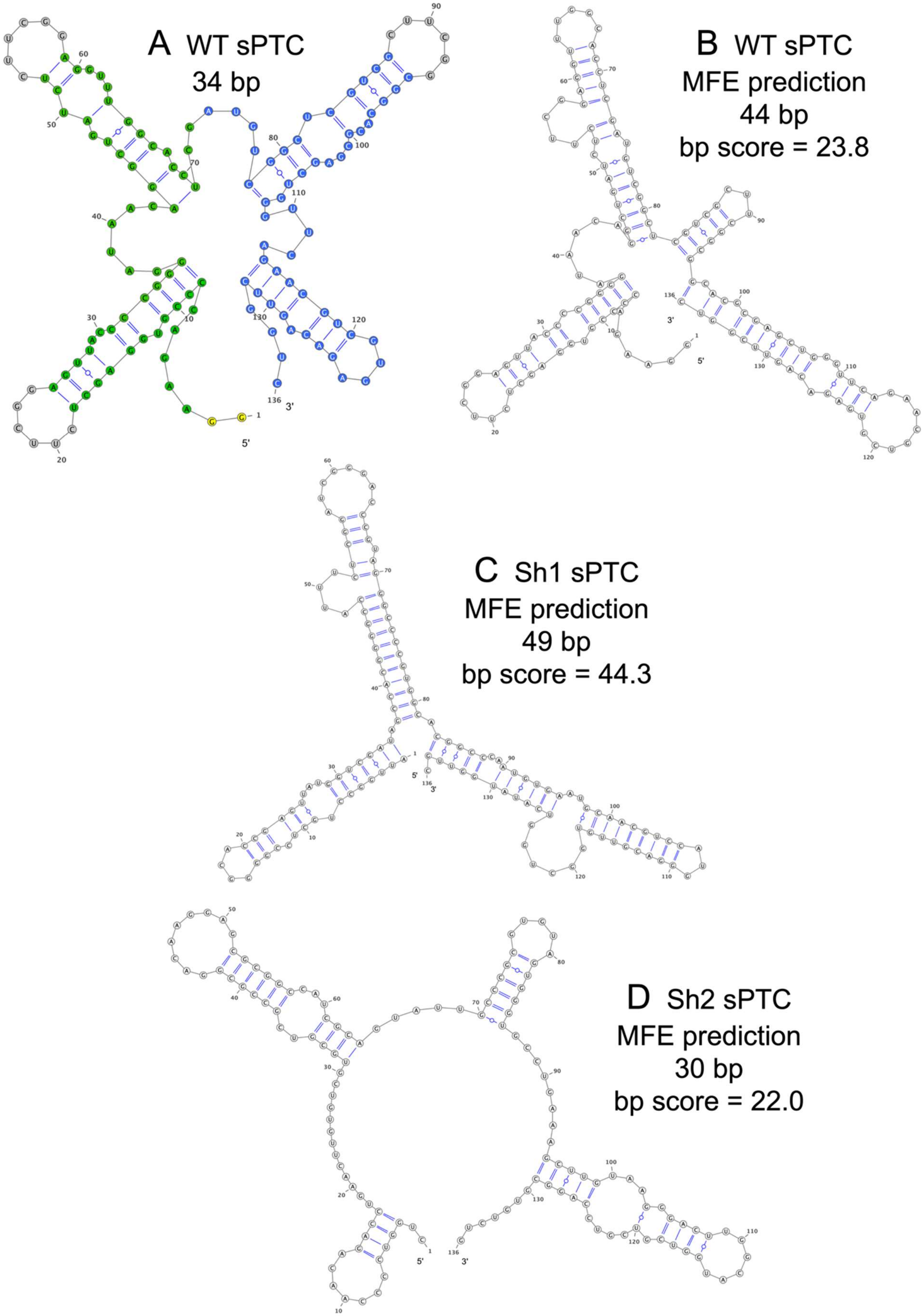
A WT sPTC
34 bp
B WT sPTC
MFE prediction
44 bp
bp score = 23.8
C Sh1 sPTC
MFE prediction
49 bp
bp score = 44.3
D Sh2 sPTC
MFE prediction
30 bp
bp score = 22.0

**Supplementary Fig. 2**. Secondary structures of WT sPTC and its two shuffled variants.
(A) Original WT sPTC based on PDB 4V51, characterised by 34 total number of base pairs (bp); blue-green colour scheme matches the two symmetrical regions depicted in **Supplementary Fig. 1**, while the additional 3 CUUCGG loops and the starter GG are also inserted and coloured in grey and yellow, respectively.
(B) NUPACK MFE prediction for the WT sPTC, characterised by 44 bp and a “bp score” (see below) of 23.8.
(C - D) NUPACK MFE prediction for the Sh1 and Sh2 sPTC, having respectively more and less structure content compared to the WT sPTC: Sh1 sPTC is characterised by 49 predicted bp and a “bp score” (see below) of 44.3, while Sh2 PTC has 30 predicted bp and a “bp score” of 22.
All predictions (B-D) were performed at the standard 1M NaCl and 37°C conditions adopted by the NUPACK algorithm. The number of base pairs (bp) in the Minimum Free Energy (MFE) is reported, together with a “bp score” computed as the sum of the probabilities that each nucleotide adopts the state of the MFE.

Even if the nominal conditions are different from the ones used in the experiments, it is guaranteed that the two variants assume different conformations, with less and more base pairs at equilibrium with respect to WT sPTC. This is due to the fact that: (i) the predicted structure of the original WT sPTC at 1 M NaCl and 37°C (B) lies in between the structures of Sh1 sPTC and Sh2 sPTC; (ii) monovalent salts and temperature have a monotonic effect on the stability of RNA.

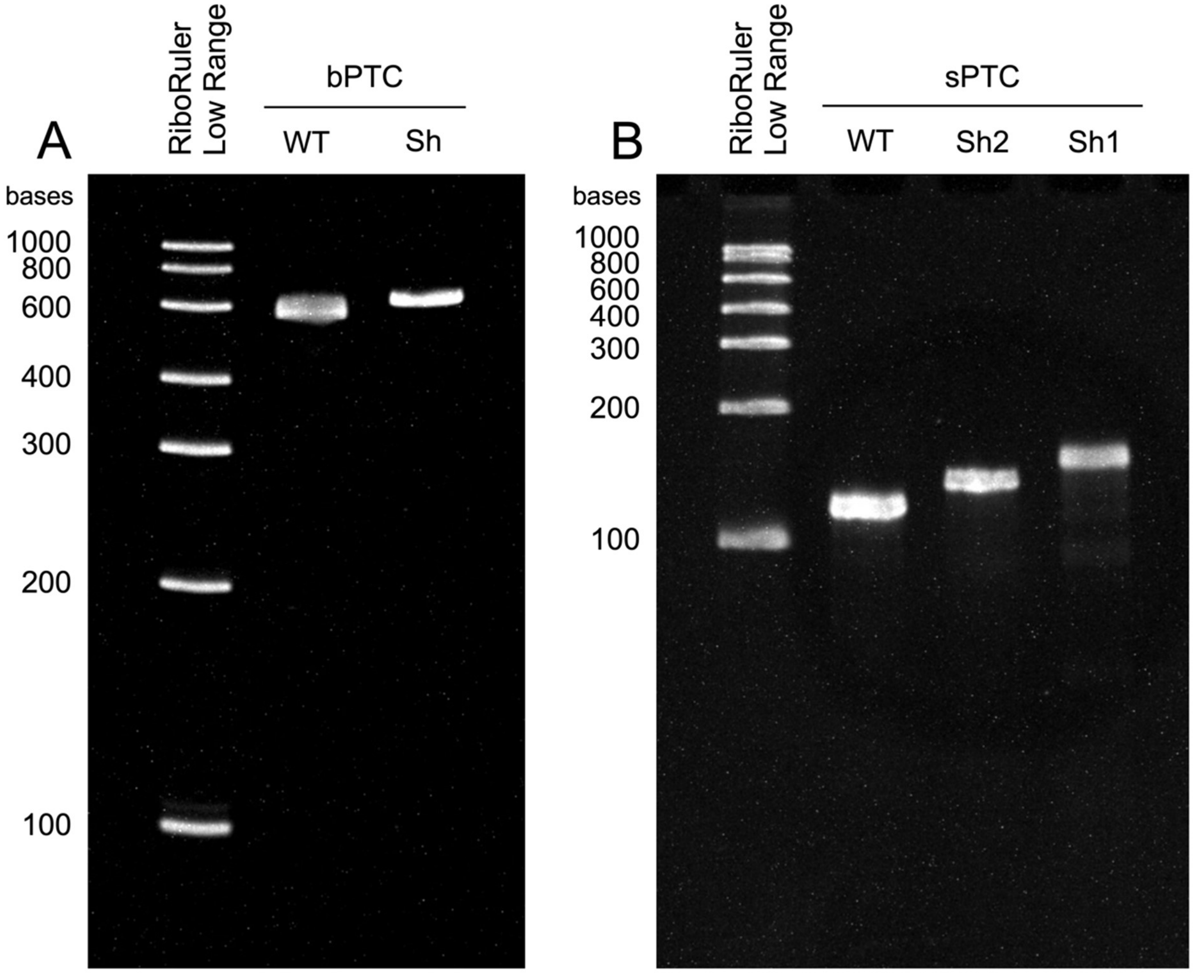


**Supplementary Fig. 3**. Denaturing gels of the RNA constructs.
(A) 5% TBE-urea gel of WT and Sh bPTC RNA constructs.
(B) 8% TBE-urea gel of WT and Sh sPTC RNA constructs.
Prior to electrophoresis, 200 ng RNA samples were mixed with RNA Gel Loading Dye (2X) (ThermoFisher Scientific) and denatured at 70°C for 10 min. RiboRuler Low Range RNA Ladder (ThermoFisher Scientific) was used as a reference. After electrophoresis, gels were soaked in TBE buffer supplemented with GelRed Nucleic Acid Gel Stain (Sigma Aldrich) for 10 min and visualised by UV transilluminator.

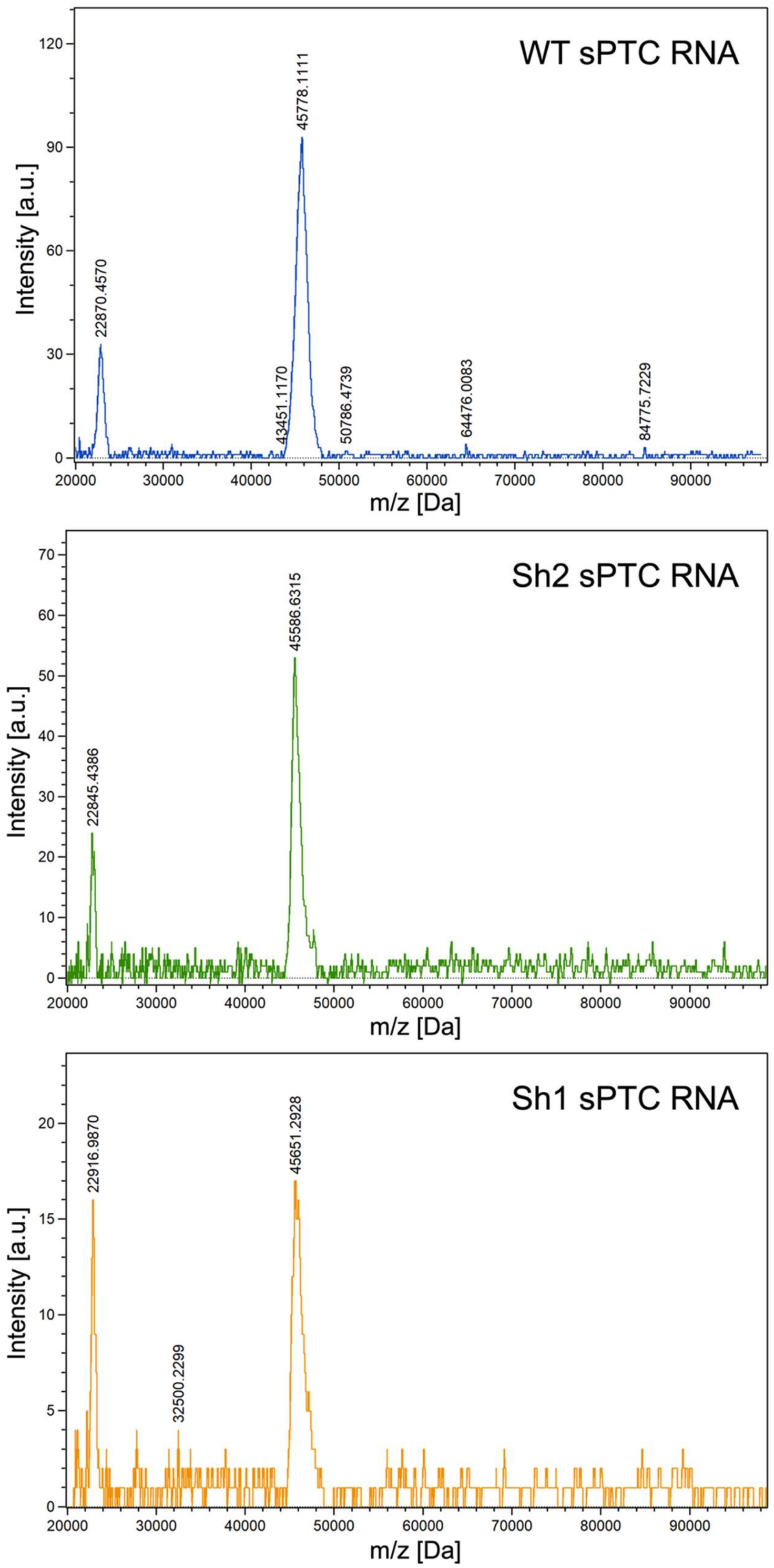


**Supplementary Fig. 4.** MALDI-TOF spectra of sPTC RNA constructs.

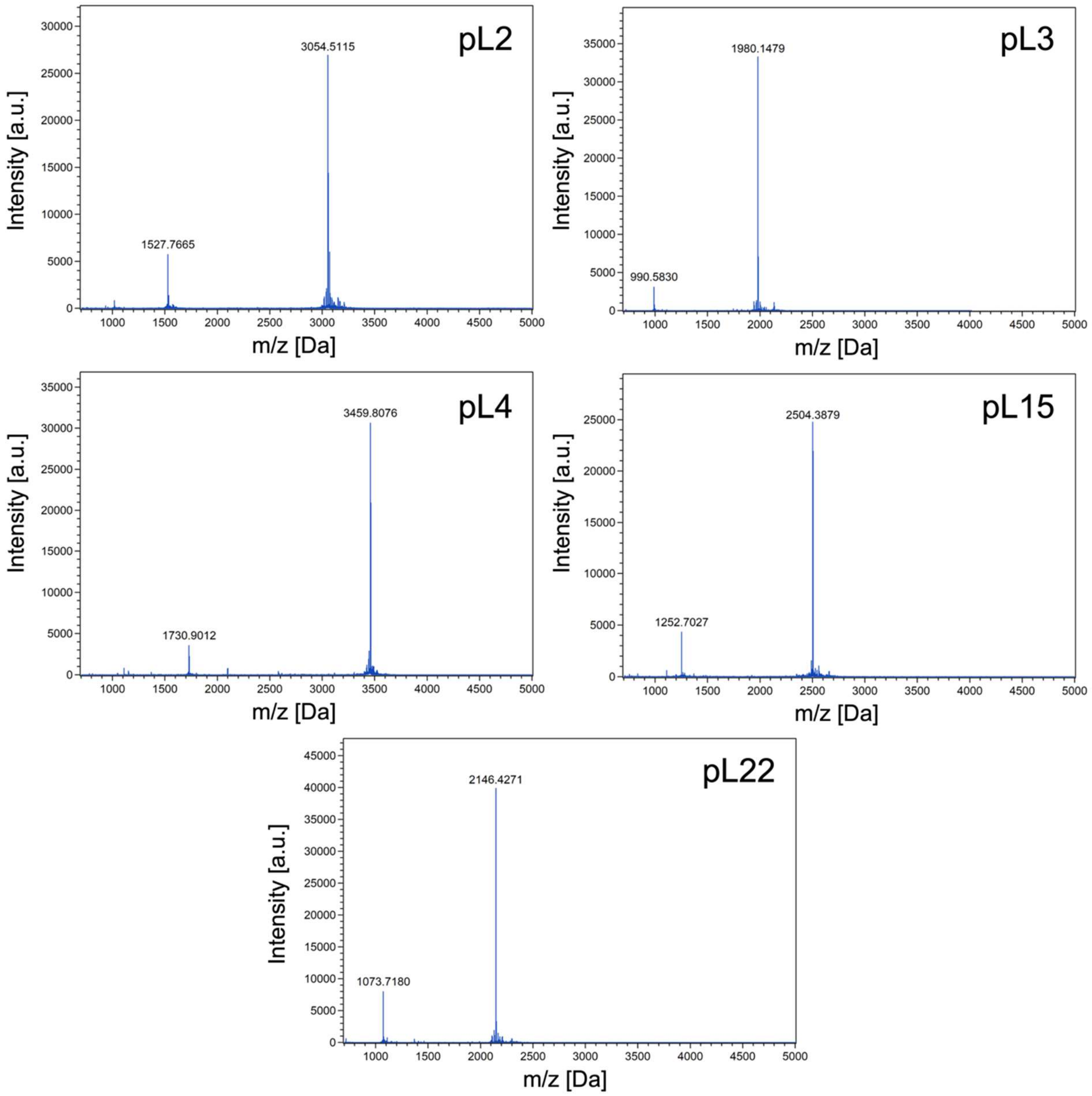


**Supplementary Fig. 5**. MALDI-TOF spectra of rPeptides (pL2, 3, 4, 15, 22).

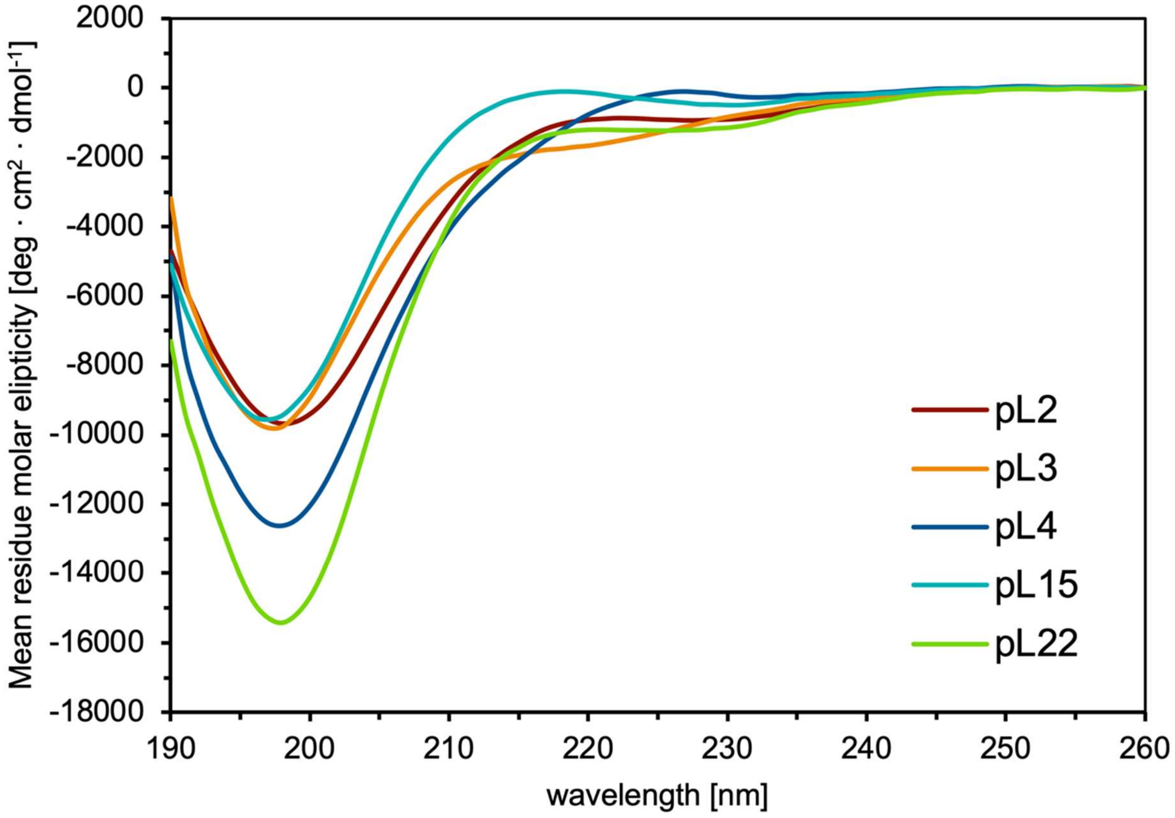


**Supplementary Fig. 6**. Far-UV CD spectra of rPeptides (pL2, 3, 4, 15, 22). The spectra were collected twice for every peptide in 10 mM Tris-HCl (pH 7.5), 1 mM KCl, 1 mM $MgCl_2$, 1 mM $CaCl_2$; the average spectra are shown.

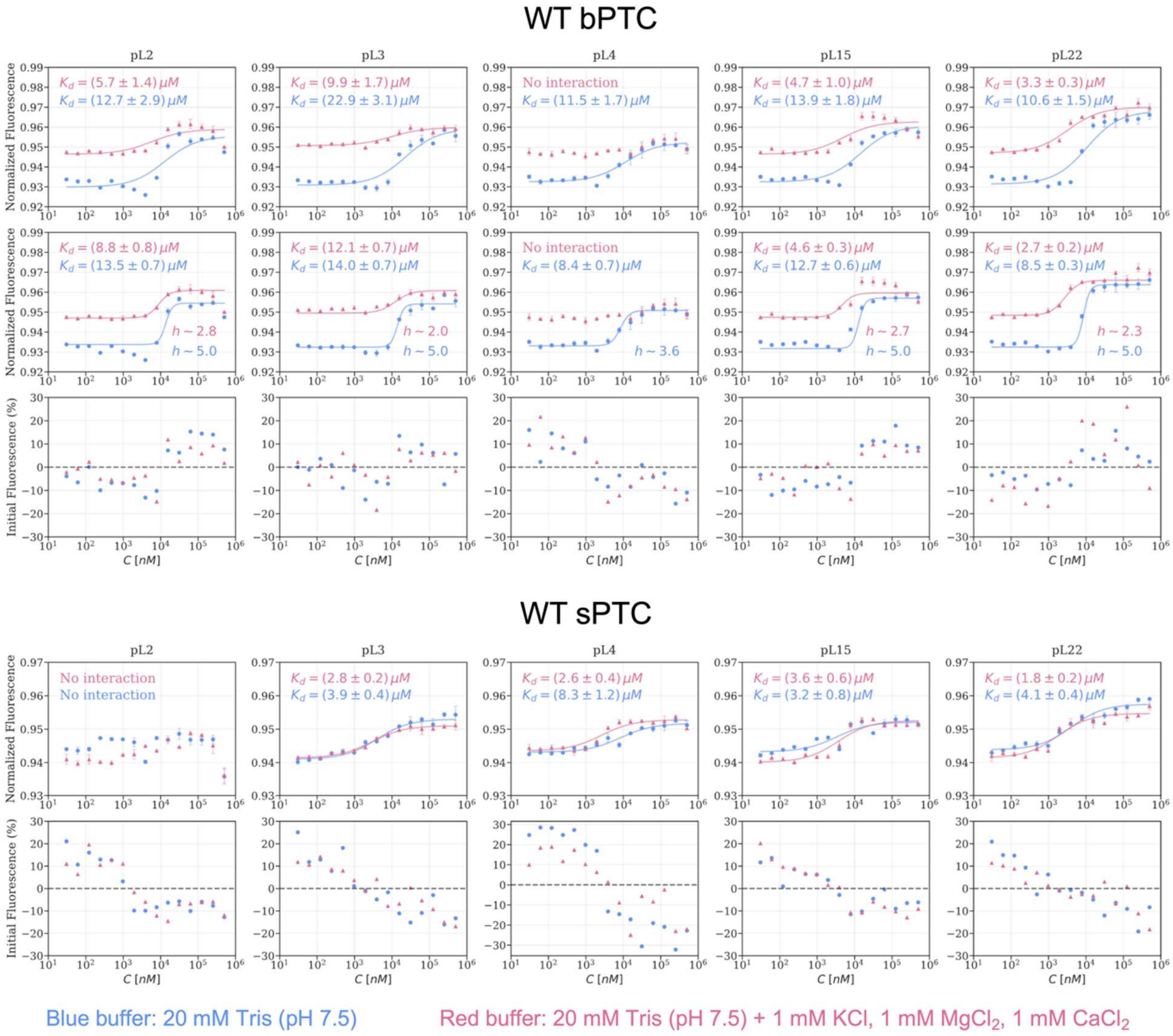


**Supplementary Fig. 7.** The binding curves (*upper rows*) and fluorescence traces (*lower rows*) for WT bPTC (*top panel*) and sPTC (*bottom panel*) with rPeptides pL2, 3, 4, 15, and 22 (*columns from left to right*), with (*red*) and without (*blue*) metal ions supplemented in the buffer. Solid curves in the first row of both panels represent the best fit to mass-action kinetics (*see eq. M below*). For WT bPTC, differences in the binding curves were observed in the presence vs absence of metal ions: the additional middle row in the WT bPTC panel shows the best fit to Hill functions (*see eq. H below*), where the approximate best Hill coefficients (denoted by *h*) were estimated.

Binding curves were analysed based on mass-action or Hill equations. sPTC curves are well fitted with a simple 1:1 binding reaction model:

$$F_n([L];K_d) = F_n^{(u)} + \frac{F_n^{(b)} - F_n^{(u)}}{2} \frac{[L] + [T_0] + K_d - \sqrt{([L] + [T_0] + K_d)^2 - 4[L][T_0]}}{[T_0]} \qquad (eq. M)$$

where $F_n$ is the normalized MST signal as a function of $[L]$, the ligand molecule concentration (pL2, 3, 4, 15 or 22, x axis in the figure above), $[T_0]$ is the fixed labelled molecule concentration (here the 50 nM WT sPTC or bPTC), $K_d$ is the dissociation constant and $F_n^{(u)}$ and $F_n^{(b)}$ are, respectively, the normalized fluorescence values of the unbounded and bounded states; binding curves were fitted against $[L]$ with free parameters $F_n^{(u)}$, $F_n^{(b)}$ and $K_d$.

On the contrary, the shape of the bPTC curves suggests binding with peptide:RNA stoichiometry significantly different from 1:1, which can be accounted for in the most simple terms by the cooperative model represented by Hill equation:

$$F_n([L]; K_d, h) = F_n^{(u)} + \frac{F_n^{(b)} - F_n^{(u)}}{1 + (K_d/[L])^h} \quad (eq.H)$$

where $[L]$, $F_n^{(u)}$, $F_n^{(b)}$ and $K_d$ have the same meaning as in $(eq.M)$, while $h$ represents the Hill exponent; binding curves were fitted against $[L]$ with free parameters $F_n^{(u)}$, $F_n^{(b)}$, $K_d$ and $h$.

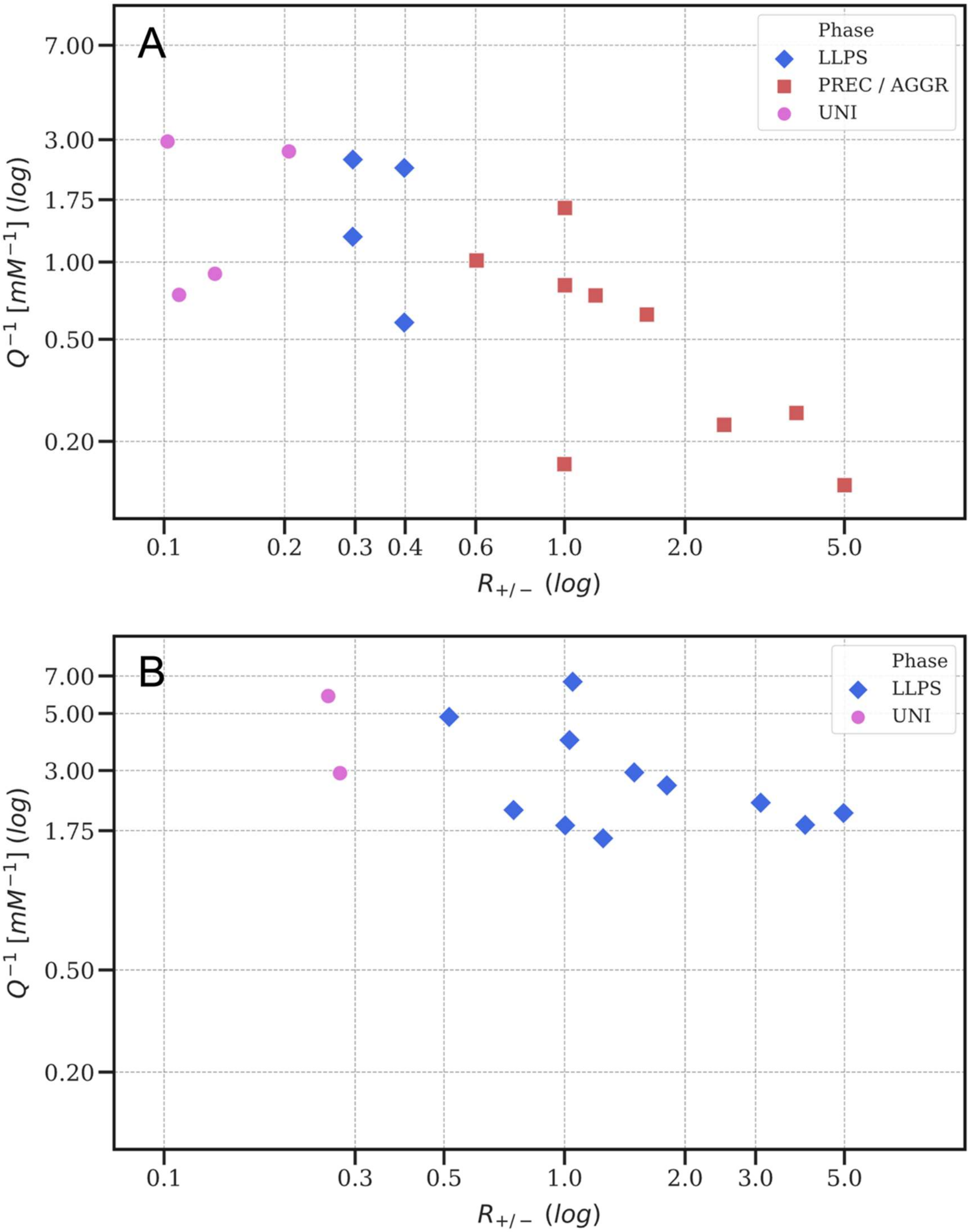


**Supplementary Fig. 8**. Phase diagrams of pL22 with WT bPTC (*panel A*) and WT sPTC (*panel B*) as a function of the stoichiometric charge ratio $R_{+/-}$ (x axis, see *eq. 1* in **Mat. & Meth., *coacervation***) and of the inverse of the molar concentration of charged units $Q^{-1}$ (y axis, see *eq. 2* in ***ibidem***). Both axes are in log scale. Different colours and markers are associated to different phases, with "UNI" indicating mixed states, "PREC/AGGR" the presence of precipitates or aggregates (**Figure 4B**) and "LLPS" the formation of liquid-like droplets (**Figure 4A & F**).

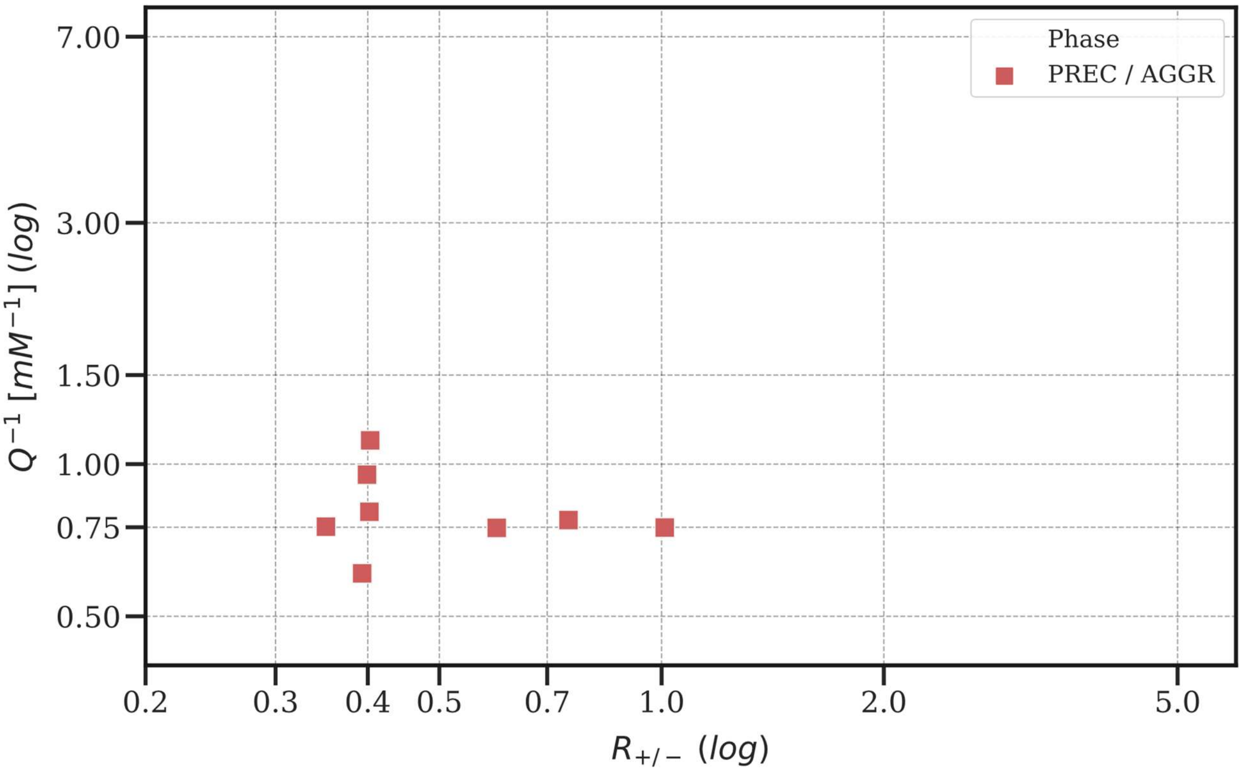


**Supplementary Fig. 9**. Phase diagram of mixtures of pLs (2, 3, 4, 15 and 22) with Sh bPTC as a function of the stoichiometric charge ratio $R_{+/-}$ (x axis, see *eq. 1* in **Mat. & Meth., *coacervation***) and of the inverse of the molar concentration of charged units $Q^{-1}$ (y axis, see *eq. 2* ***ibidem***). Both axes are in log scale. Every sample examined contained some precipitates or aggregates (**Figure 4B**), thus the phase is indicated with "PREC/AGGR", with the same colour and marker code as in **Figure 4** and in **Supplementary Fig. 8**.

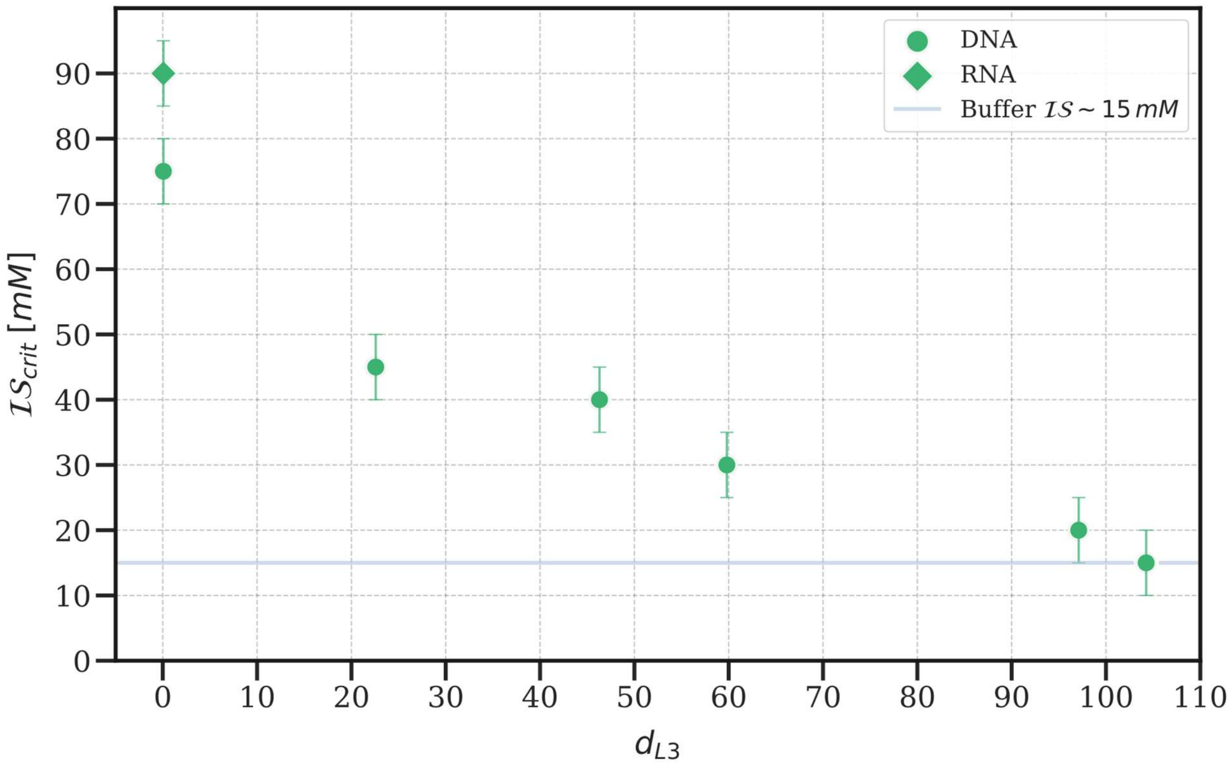


**Supplementary Fig. 10**. Scatter plot of the critical ionic strength (y axis) as a function of the distance (x axis) between the composition of the pL3 environment and that of several ssDNA unstructured oligomers and one RNA strand (see the legend and the "RNA" and "DNA" tags).

The critical ionic strength, measured at the same fixed DNA/RNA and peptide concentration and at ambient temperature for all the peptide-nucleic acid pairs, is defined as the highest NaCl concentration at which coacervation is still observed under a bright field microscopy setting. The distance $d_{L3}$ is defined using the euclidean metric as:

$$d_{L3} = \sqrt{(A - A_{L3})^2 + (C - C_{L3})^2 + (G - G_{L3})^2 + (T - T_{L3})^2}$$

where A, C, G, T indicate the nucleobases percentage composition of the oligomers, while the same letters with "L3" subscript refers to the percentage composition of the pL3 *T. Thermophilus* rRNA environment, measured directly locating the 5 Angstrom closer residues to pL3 in PDB 4V51. Error bars account for a 10 mM experimental uncertainty on the measured critical ionic strength, while the grey horizontal line indicates the approximate buffer ionic strength (20 mM Tris, pH 7.6), which constitutes the experimental lower limit.

A

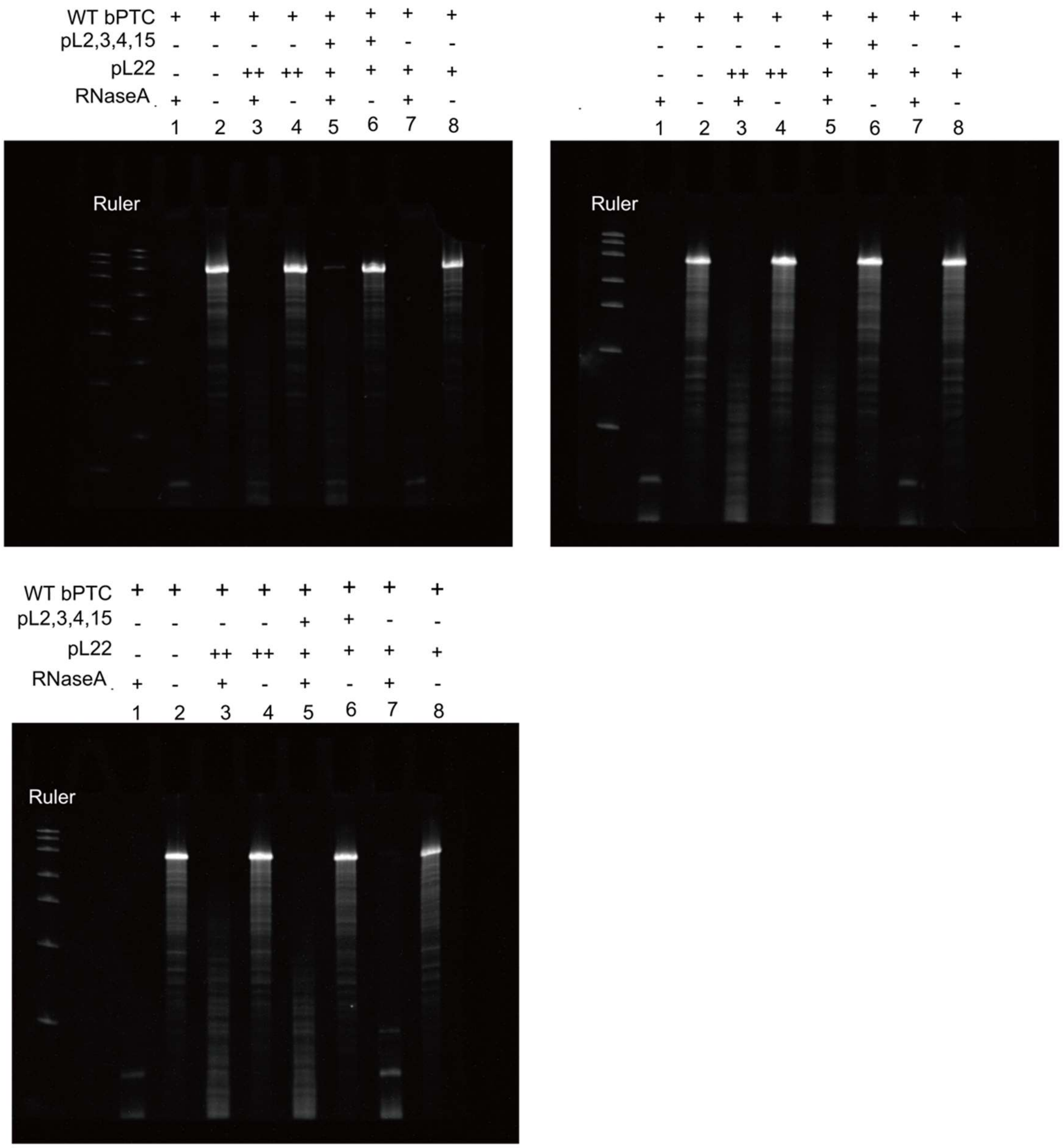
WT bPTC + + + + + + + +
pL2,3,4,15 - - - - + + - -
pL22 - - ++ ++ + + + +
RNaseA + - + - + - + -
1 2 3 4 5 6 7 8
Ruler
+ + + + + + + +
- - - - + + - -
- - ++ ++ + + + +
+ - + - + - + -
1 2 3 4 5 6 7 8
Ruler
WT bPTC + + + + + + + +
pL2,3,4,15 - - - - + + - -
pL22 - - ++ ++ + + + +
RNaseA + - + - + - + -
1 2 3 4 5 6 7 8
Ruler

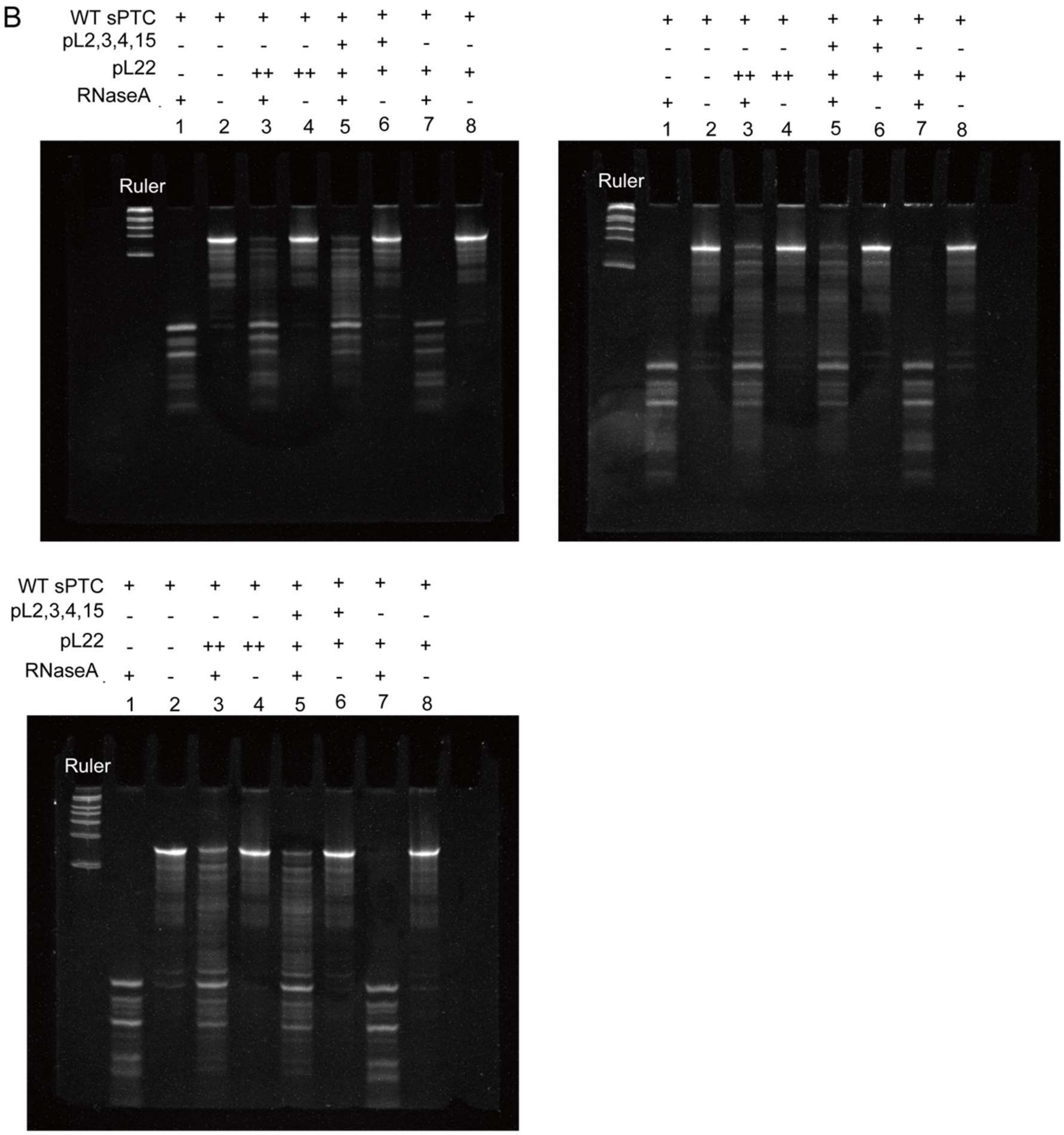


**Supplementary Fig. 11**. Full size Urea-gel analysis of the WT bPTC (*A*) and sPTC (*B*) RNaseA stability (see **Figure 5**). “Ruler” corresponds to RiboRuler ™ Low Range RNA ladder ready to use, Thermo Fisher Scientific.

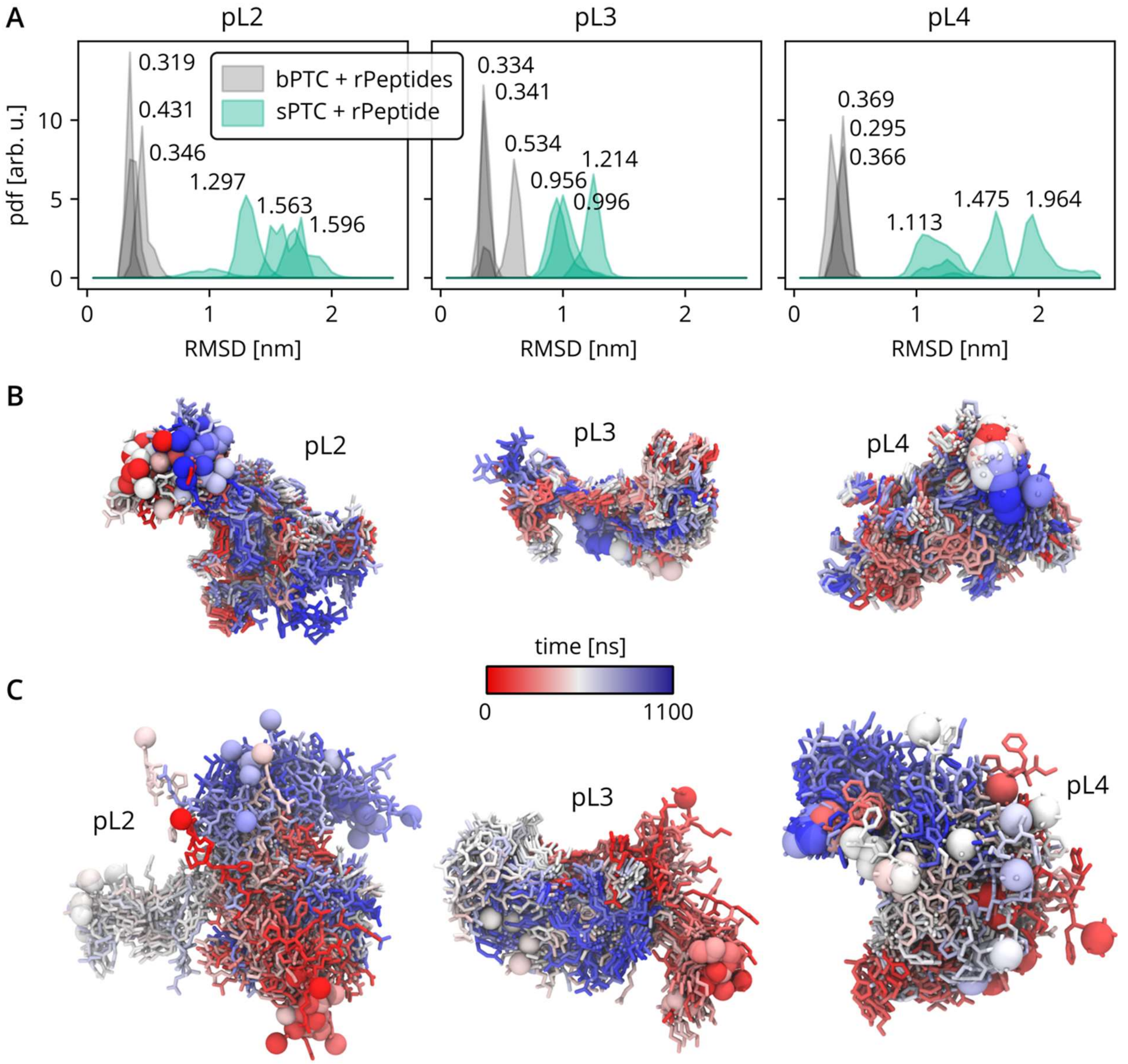


**Supplementary Fig. 12.** Analysis of selected rPeptides (pL2, pL3 and pL4).

(A) The root-mean-square deviation (RMSD) of rPeptides with respect to their initial position after least-square superposition of the largest common rRNA element obtained from MD simulations of WT bPTC containing all rPeptides (*grey*) or WT sPTC with one rPeptide (*green*). The mean values of the probability distribution functions (pdfs) are denoted.

(B) Representative rPeptide conformations on the bPTC surface (*not shown*), color-coded according to the simulation time from red through white to blue. The spheres represent the C-terminal $C\alpha$ atoms.

(C) Same as (*B*) but for the MD simulations of the sPTC + rPeptide.
